# An intracellular hydrophobic nexus is critical for slow deactivation in hERG channels

**DOI:** 10.1101/2023.03.04.531007

**Authors:** Whitney A. Stevens-Sostre, Lisandra Flores-Aldama, Daniel Bustos, Jin Li, Lucie Delemotte, Gail A. Robertson

**Author notes:** These authors equally contributed to the manuscript (Whitney A. Stevens-Sostre, Lisandra Flores-Aldama). Correspondence to Gail Robertson.

## Abstract

Slow deactivation is a critical biophysical and physiological property of voltage-gated K^+^ channels encoded by the *human Ether-à-go-go-Related Gene* (*hERG*). hERG channel deactivation is modulated by interactions between intracellular N-terminal Per-Arnt-Sim (PAS) and C-terminal cyclic nucleotide-binding homology (CNBh) domains. The PAS domain is multipartite, comprising a globular domain (residues 26-136) and an N-terminal PAS-cap that is further subdivided into an initial unstructured segment (residues 1-12) and an amphipathic α-helical region (residues 13-25). Disruption of interactions within the PAS-CNBhD complex accelerates deactivation kinetics and impairs the slow repolarization of the cardiac action potential. Similarly, deletion of the PAS-cap unstructured segment, modeled as a "foot-in-the-door,” accelerates channel closing. Here, we tested the hypothesis that a three-dimensional hydrophobic nexus, comprising residues in the PAS-cap α-helix, the globular PAS, and CNBh domains, plays a crucial role in slow deactivation. Utilizing structure-directed mutagenesis, electrophysiology, and molecular dynamics simulations, we provide evidence that altering hydrophobicity at this interface accelerates deactivation kinetics and causes a rearrangement in the network of residues comprising the hydrophobic nexus. Moreover, the PAS-cap unstructured segment disengages from the gating machinery. We propose that the nexus hydrophobicity positions the PAS-cap α-helix and promotes engagement of the upstream PAS-cap unstructured segment with the gating machinery, resulting in slow channel closing. Interestingly, several long QT syndrome disease mutations lie at this interface, underscoring the importance of the hydrophobic nexus and the novel role of the PAS-cap α-helix in hERG gating.

## Introduction

The voltage-gated K^+^ channel Kv11.1, encoded by the *human Ether-à-go-go (EAG)-Related Gene* (*hERG*), or *KCNH2*, contributes to several physiological processes, including cardiac repolarization (Sanguinetti et al., 1995; Trudeau et al., 1995), neuronal excitability (Sacco et al., 2003; Furlan et al., 2007; Huffaker et al., 2009), and cellular proliferation (Crociani et al., 2003). These channels, commonly referred to as human ERG (hERG) channels, give rise to the rapid delayed-rectifier K^+^ current (I_Kr_) critical for the repolarization of the ventricular action potential (Sanguinetti et al., 1995; Trudeau et al., 1995). hERG slow deactivation is a key biophysical and physiological property that influences the shape of I_Kr_ currents (Sanguinetti et al., 1995; Zhou et al., 1998). Unintended pharmacological block of hERG channels causes acquired long QT syndrome (LQTS) (Roden, 2006; Vandenberg et al., 2012), while congenital mutations in the *KCNH2* gene cause type 2 LQTS (Curran et al., 1995; Gustina and Trudeau, 2012) and are implicated in schizophrenia (Huffaker et al., 2009; Atalar et al., 2010; Apud et al., 2012; Hashimoto et al., 2013), epilepsy (Johnson et al., 2009; Zamorano- Leon et al., 2012), and cancer (Bianchi et al., 1998; Pointer et al., 2017). Identifying the molecular determinants underlying proper hERG channel function is, therefore, crucial to understanding how life-altering diseases can result from ion channel dysfunction.

Tetrameric ERG and EAG channels, members of the KCNH superfamily of voltage-gated K^+^ channels, share structural similarities in their intracellular amino (N) and carboxyl (C) termini (Whicher and MacKinnon, 2016; Wang and MacKinnon, 2017). Distinctive features of KCNH channels are the cytosolic N-terminal Per-Arnt-Sim (PAS) (Morais Cabral et al., 1998) and C-terminal cyclic nucleotide binding homology (CNBh) domains (Guy et al., 1991; Marques-Carvalho et al., 2012) (**Fig. 1**). The PAS domain (residues 1-135) comprises the N-terminal PAS cap (residues 1-25) and globular PAS (residues 26-135) domains (Morais Cabral et al., 1998). Cryogenic electron microscopy (Cryo-EM) and nuclear magnetic resonance studies reveal that the PAS-cap is further divided into an unstructured region (residues 1-12) and an amphipathic α-helix (residues 13-25) (Gayen et al., 2011; Muskett et al., 2011; Ng et al., 2011; Wang and MacKinnon, 2017). The CNBh domain is structurally similar to cyclic nucleotide-binding domains of hyperpolarization-activated cyclic nucleotide–gated (HCN) and cyclic nucleotide-gated (CNG) channels (Guy et al., 1991) but does not bind cyclic nucleotides (Robertson et al., 1996; Brelidze et al., 2009). The globular PAS and CNBh domains interact to form an inter-subunit intracellular “gating ring” with clear modulatory roles in KCNH channel gating (Gustina and Trudeau, 2011; Haitin et al., 2013; Morais-Cabral and Robertson, 2015; Robertson and Morais-Cabral, 2020; Harley et al., 2021). The structural arrangement of both the globular PAS and CNBh domains are highly conserved, and their interacting interface is a hotspot for disease-related mutations (Haitin et al., 2013; Foo et al., 2019), underscoring their importance in survival across a wide phylogenetic spectrum.

**Fig. 1.**
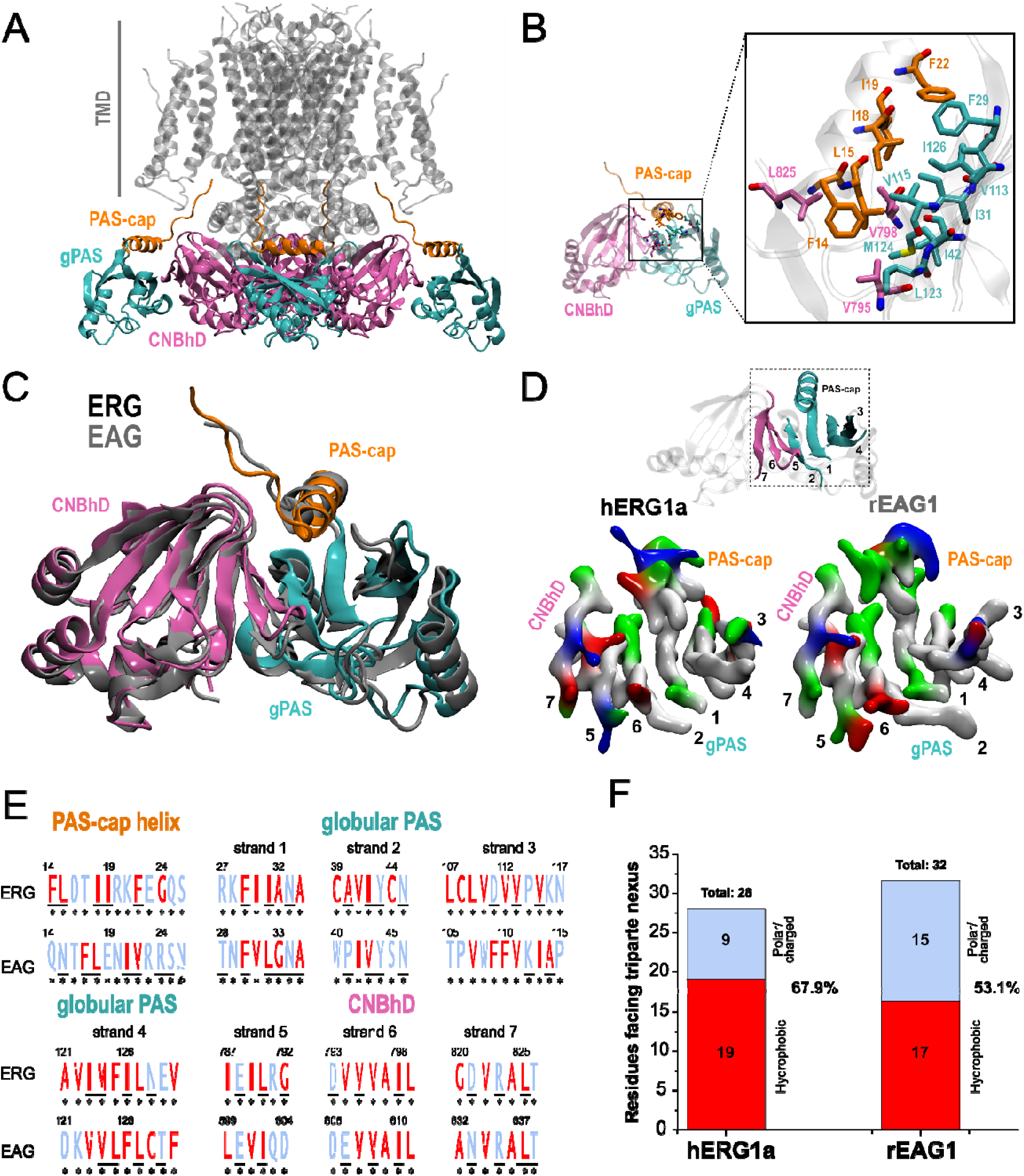
Structural analysis reveals a tripartite hydrophobic nexus between hERG1a intracellular domains. (**A**) Cryo-EM structure of hERG1a tetramer (PDB entry: 5VA1) highlighting the intracellular PAS-cap (orange), globular PAS (gPAS) (cyan), cyclic nucleotide-binding homology domain (CNBhD) (pink), and transmembrane domains (TMD) (grey). (**B**) The intracellular PAS-cap α-helix, gPAS, and CNBhD form a hydrophobic nexus whose constituent residues are highlighted in the inset. (**C**) Structural alignment between the hERG1a (PDB entry: 5VA1) and rEAG1 (PDB entry: 5K7L) intracellular N-terminal PAS and C-terminal CNBh domains. (**D**) Color-coded hydropathy of residues facing the proposed hydrophobic nexus. Amino acids are colored gray for hydrophobic, green for hydrophilic, blue for basic, and red for acidic properties. Strands were numbered according to sequence, and their positioning within the interface is shown in the inset. (**E**) Sequence identity (by letter size) of amino acids located at motifs constituting the proposed hydrophobic nexus in mammalian ERG1a and EAG1 ion channels. Hydrophobic and hydrophilic residues are colored red and blue, respectively. Residues facing the PAS-cap α-helix/ globular PAS/CNBhD interface are underlined. The number of the strands corresponds to numeration in (D). (**F**) Hydropathy of amino acid side chains facing the tripartite interface. Calculations were made according to alignment in (D) and available structures of hERG1a (PDB entry: 5VA1) and rEAG1 (PDB entry: 5K7L). Multiple sequence alignment in Supplemental Fig. S1 was used to calculate the percentage of amino acid conservation (see Materials and Methods section). Asterisks (*) indicate an amino acid sequence identity of 100%.

Previous studies showed that deleting the entire N-terminal region (Δ2-354) accelerates hERG deactivation, a phenotype also observed after deleting only the N-terminal PAS-cap disordered region or tip (residues 2-16) (Schonherr and Heinemann, 1996; Morais Cabral et al., 1998; Wang et al., 1998). Reintroduction of a peptide corresponding to the deleted 16 amino acids is sufficient to reconstitute approximately 50% of the slow deactivation to the N-terminal truncated channels (Wang et al., 2000). This work and studies by others led to a “foot-in-the-door” model by which closing normally occurs only after the PAS-cap tip disengages from the gating machinery and a critical energetic barrier is removed. This model is supported by other functional studies and cryo-EM ERG structures showing the PAS-cap tip in close apposition with the S4-S5 linker at the bottom of the voltage-sensor domain (VSD) in its activated position, leading to the hypothesis that the PAS-cap tip exits this site when the VSD moves to its resting position, permitting the channel to close (Wang et al., 1998; de la Pena et al., 2011; Fernandez-Trillo et al., 2011; Ng et al., 2012; Hull et al., 2014; Wang and MacKinnon, 2017). In addition, “hydrophobic patches” previously identified on the surfaces of the globular PAS (Morais Cabral et al., 1998) and CNBh domains (Al-Owais et al., 2009) were suggested to modulate hERG deactivation (Morais Cabral et al., 1998). Nevertheless, the allosteric pathway communicating the PAS-cap tip with these cytosolic regions during deactivation remains unknown.

Here, upon inspection of the cryo-EM structure of the hERG channel, we identified a hydrophobic interface involving three discrete structural domains: the PAS-cap α-helix, the globular PAS domain, and the CNBh domain. We tested the hypothesis that this nexus of three modular regions plays an active role in slow deactivation. Site-directed mutagenesis, electrophysiology, and molecular dynamics simulations (MDS) analyses show that reducing the hydrophobicity within this nexus consistently accelerates deactivation and evokes a retraction of the PAS-cap tip from the gating machinery.

## Results

### A hydrophobic interface among the PAS-cap α-helix, globular PAS, and CNBh domains is characteristic of mammalian ERG channels

Inspection of the cryo-EM structure of the tetrameric hERG channel (PDB:5VA1) reveals the PAS-cap α-helix (residues 13-25) located at an interface between the globular PAS (residues 26-135) and CNBh domains (residues 736-863), while the PAS-cap unstructured segment (residues 1-12) lies apposed to the S4-S5 linker in the transmembrane region (**Fig. 1A-B**) (Wang and MacKinnon, 2017). **Fig. 1B** reveals a “hydrophobic patch” along the amphipathic PAS-cap α-helix comprising residues F14, L15, I18, I19, and F22, in orange. These residues are within 4.5 Å of hydrophobic amino acids at the interface between the globular PAS and CNBh domains, in teal and pink, respectively. Together, these hydrophobic residues form a three-dimensional nexus of three constituent modules.

Searching for structural underpinnings of the slow deactivation in ERG that are not shared by the structurally similar EAG1 channel (Robertson et al., 1996), we compared the overall structure and amino acid conservation of the motifs contributing to the hydrophobic nexus in the two channels. Overall structural homology between the PAS-cap, globular PAS, and CNBh domains of hERG and the closely related rat EAG1 channel (rEAG1) (Wang nad MacKinnin, 2017; Whicher and MacKinnon, 2016) is apparent by overlapping the two structures (**Fig. 1C**). Color-coded hydropathy of residues at the tripartite interface in each channel revealed a higher density and closer proximity of hydrophobic residue side chains (in gray) in hERG relative to rEAG1, suggesting tighter hydrophobic interactions (**Fig. 1D**). Multiple sequence alignment of 20 mammalian ERG (ERG1a isoform) and EAG1 (long isoform) sequences showed near 100% identity in the motifs contributing to the tripartite nexus within the respective groups (**Fig. S1**; **Fig. 1E**), reflecting the importance of the interface for both channel types. However, when comparing the sequences between mammalian EAG1 and ERG1a channels, differences are evident (**Fig. 1E**). Hydropathy analysis of residues at this interface showed a greater percentage of hydrophobic residues in mammalian ERG1a (68%) compared to EAG1 channels (53%) (residues in red in **Fig. 1E-F**). These primary and tertiary structural differences suggest that mammalian ERG1a channels evolved towards a unique hydrophobic nexus and lead to the hypothesis that this tripartite interface is important in establishing slow deactivation, a key defining characteristic of gating in hERG channels.

### A hydrophobic patch within the PAS-cap α-helix contributes to slow deactivation kinetics

To test this hypothesis and determine the role of hydrophobicity within the hERG three-dimensional nexus, we initially mutated hydrophobic residues on the PAS-cap α-helix to serine (**Fig. 2A**), an average-sized hydrophilic amino acid. Additionally, we performed deletions of the hERG PAS-cap unstructured region (Δ2-12), PAS-cap α-helix (Δ13-25), or entire PAS-cap domain (Δ2-25) to establish and compare their respective hypomorphic phenotypes. We determined the effects of deletions and serine substitutions on deactivation kinetics using two-electrode voltage clamp (TEVC) in *Xenopus laevis* oocytes expressing WT or mutant hERG channels (**Fig. 2B**). Calculation of the weights of two-exponential fits to the currents showed the fast component contributes ∼90% of the deactivating current at the test potential applied (i.e., –120 mV) for all mutants (**Fig. S2** and **Table S1**, see Materials and Methods). Thus, for simplicity, only the fast time constants of deactivation ( ) are reported in this study.

**Fig. 2.**
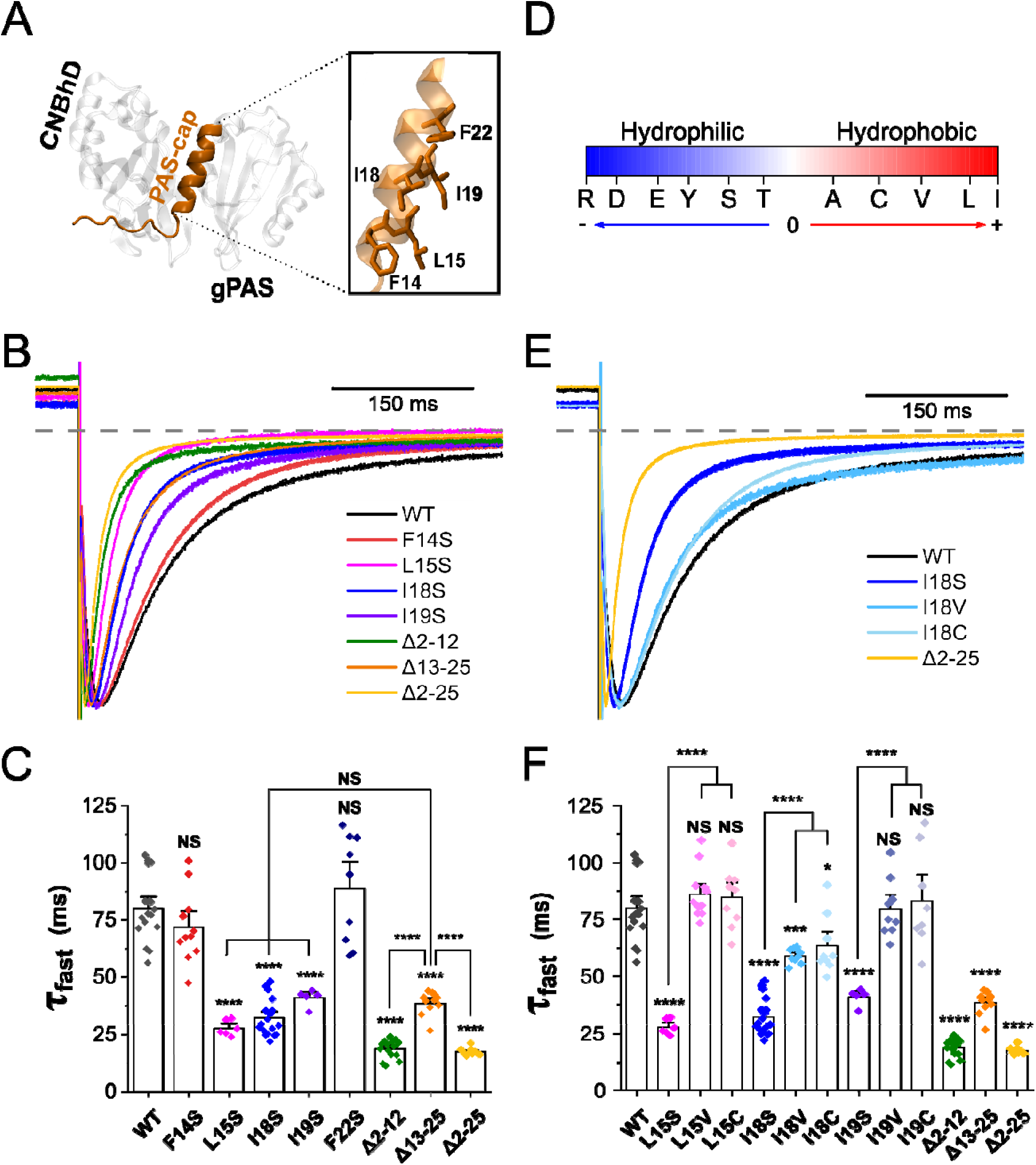
Hydrophobicity of PAS-cap α-helix residues determines slow deactivation in hERG1a channels. (**A**) Representation of one of the four PAS-CNBhD complexes forming the “gating ring” in hERG1a channels (PDB entry: 5VA1). The positioning of the PAS-cap domain is highlighted in orange. The inset highlights the residues at the PAS- cap α-helix contributing to the proposed hydrophobic nexus. (**B**) Scaled tail currents of WT (wild-type) and PAS-cap F14S, L15S, I18S, I19S, Δ2-12, Δ13-25, and Δ2-25 mutant channels. (**C**) Time constant of the fast component of deactivation in WT and PAS-cap mutant channels. (**D**) Scheme representing the hydropathy of amino acids relevant to our study according to Goldman-Engelman-Steitz hydropathy scale. (**E**) Scaled tail currents of WT and PAS-cap I18S, I18V, I18C, and Δ2-25 mutant channels. (**F**) Time constant of the fast component of deactivation in WT and PAS-cap L15S, L15V, L15C, I18S, I18V, I18C, I19S, I19V, I19C, Δ2-12, Δ13-25, and Δ2-25 mutant channels. Tail currents in (**B**) and (**E**) were evoked at -120 mV after a step to +60 mV. Dotted line indicates zero current level. Data in (**C**) and (**F**) are mean ± SEM. (****, P < 0.0001; ***, P < 0.001; *, P < 0.05; NS: not significant, P > 0.05).

Analysis of the truncated channels showed a faster current decay when compared to WT hERG, suggesting both the disordered region (Δ2-12) and the PAS-cap α-helix (Δ13-25) contribute to slow deactivation gating (**Fig. 2B-C**, **Fig. S2B-E**, and **Table S2**), as previously reported (Schonherr and Heinemann, 1996; Wang et al., 1998; Ng et al., 2011). Interestingly, deleting the PAS-cap α-helix (Δ13-25) yields slower deactivation relative to deletions encompassing the extreme N-terminal 12 or 16 residues (**Fig. 2C**). Because deletion of the N-terminal PAS-cap residues (Δ2-12) mimics loss of the entire N-terminus (Wang et al., 1998), we infer that the Δ13-25 deletion results in a less extreme phenotype with the PAS-cap unstructured segment engaging the gating machinery less effectively than in the WT channel.

PAS-cap mutations L15S, I18S, and I19S on the hydrophobic patch of the helix accelerated deactivation kinetics, with that were not statistically different from those of the Δ13-25 deletion but statistically slower than deletions comprising the N-terminal disordered region (Δ2-12 and Δ2-25). Serine substitutions of flanking residues F14 and F22 did not accelerate deactivation and thus represent the functional boundaries of the hydrophobic patch actively engaged in the nexus (**Fig. 2B-C** and **Fig. S2F-G**). Steady-state voltage dependence of activation did not differ among the mutant, truncated, or WT channels, with the exception of the Δ2-12 (i.e., PAS-cap disordered region), which produced a right-ward shift in the half–maximal voltage (V_1/2_) of the current-voltage relation, as has been previously reported (Ng et al., 2011) (**Fig. S2** and **Table S3**). These results show that individual L15S, I18S, and I19S mutations reproduce the Δ13-25 phenotype and suggest they disrupt contributions of the PAS-cap α-helix (residues 13-25) to the hydrophobic nexus and deactivation allosteric pathway.

To further test the importance of hydrophobicity of the PAS-cap α-helix in deactivation gating, we substituted residues L15, I18, and I19 with residues predicted to have intermediate hydrophobicity per the Goldman-Engelman-Steitz hydropathy scale (Zviling et al., 2005) (**Fig. 2D)**. To avoid steric hindrance effects, we selected valine and cysteine, which have similar size to serine in mutants and leucine/isoleucine in WT channels, respectively. Substitutions to either valine or cysteine at residues L15 and I19 showed no changes in deactivation kinetics compared to WT, indicating that a slight reduction of hydrophobicity at either residue was insufficient to disrupt interactions of the PAS-cap α-helix. In contrast, I18 substitutions to valine or cysteine accelerated deactivation kinetics to a level intermediate to those of WT and the serine mutant, suggesting a more critical dependence on the hydrophobicity at that residue (**Fig. 2E-F**). A quantitatively similar result was reported for I19A, consistent with hydrophobicity playing a key role in this region (Ng et al., 2011). These results indicate that the PAS-cap α-helix contributes to hERG slow deactivation kinetics via hydrophobic interactions of a patch comprising L15, I18, and I19 and delimited by the flanking residues F14 and F22. I18, in the center of the region and most sensitive to perturbation, may play a pivotal role.

### Serine substitutions do not cause structural perturbations to the PAS-cap α-helix in molecular dynamics simulations

Serine substitutions can disrupt α-helical conformations (Ballesteros et al., 2000) and a previous study reported that the introduction of multiple glycine and serine residues to the PAS-cap α-helix disrupts its secondary structure (Ng et al., 2011). To discern between effects on local hydrophobicity versus loss of α-helix secondary structure, we performed short (300 ns) fully atomistic molecular dynamics simulations (MDS) utilizing the tetrameric hERG cryo-EM structure (PDB: 5VA1) and *in silico* generated F14S, L15S, I18S, and I19S mutant channels (**Fig. 3 and Fig. S3**). Data for the analysis was captured after the MDS equilibrium was reached. This equilibrium was defined as the point where the protein root-mean-square deviation (RMSD) stabilized and corresponds to 125 ns; therefore, 175 ns of the MDS were analyzed, as indicated by the horizontal lines in **Fig. S3**. Secondary structure analysis of WT hERG PAS-cap α-helix showed that residues 15-25 maintained a stable α-helical structure throughout the simulation time (**Fig. 3A-B**). Similar results were observed for the F14S mutant, which mimicked the WT channel slow deactivation in our functional analyses. When compared to slow deactivating WT and F14S channels, fast deactivating mutants L15S and I19S showed no changes in α-helix conformation throughout the simulation time (**Fig. 3D-F**). In contrast, I18S showed disruption of secondary structure at distal regions upstream (residue 15) and downstream (residues 23-25) from the hydrophobic patch in the second half of the simulation (**Fig. 3D-F**). Nonetheless, most of the PAS-cap α-helix hydrophobic residues contributing to the three-dimensional nexus retained α-helical structure throughout the I18S mutant simulation time. These results suggest that the secondary structure of the PAS-cap α-helix is conserved in most of the mutants and, consequently, faster deactivation kinetics observed in functional analyses arise from perturbation of local hydrophobicity. A special case is the I18S mutant, in which disruption of both hydrophobicity and α-helix secondary structure may contribute to accelerated current decay.

**Fig. 3.**
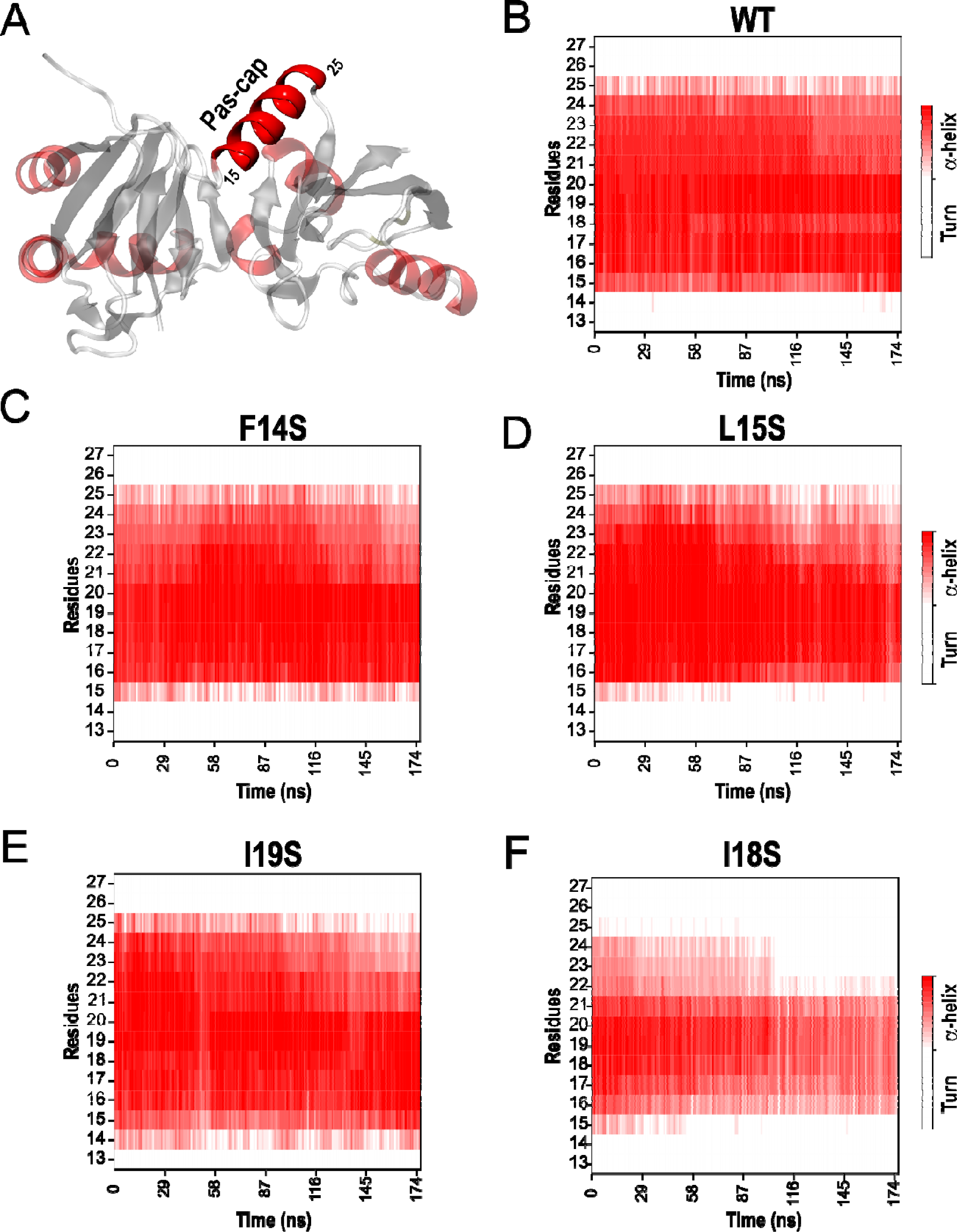
Analysis of the secondary structure of the PAS-cap α-helix in WT and serine mutant hERG1a channels throughout the equilibrium simulation time. (**A**) Representation of the secondary structure of the PAS-CNBhD complex in WT hERG1a tetramers (PDB entry: 5VA1). Color represents different secondary conformations; turn in white, β strand in gray, α-helix in red. Quantification of the secondary structure of the PAS-cap α-helix for the slow deactivating WT (**B**) and F14S (**C**) mutant, and fast deactivating mutants L15S (**D**), I19S (**E**), and I18S (**F**). Represented values are the average of the helix secondary structure in each of the four subunits in three MDS replicates, producing a total of 12 replicates.

### Residues in globular PAS and CNBh domains contribute to the hydrophobic nexus mediating slow deactivation kinetics

Given the role of hydrophobic amino acids in the PAS-cap α-helix in gating, we next studied whether proximal hydrophobic residues in the globular PAS and CNBh domains indicated in the cryo-EM structure also have an active role in deactivation through the structurally defined nexus (**Fig. 4**). We first identified residues located at the globular PAS and CNBh domains within 4.5 Å of PAS-cap α-helix hydrophobic residues L15, I18 and I19. Our analysis revealed eight hydrophobic residues on the globular PAS domain and three on the CNBh domain (**Fig. 4A**; residues L15, I18 and I19 are not shown for visual clarity). Serine substitutions at each of these residues accelerated deactivation, except for M124S, which did not show statistical differences compared to the WT channel (**Fig. 4B** and **Table S4**). Within the globular PAS domain, F29S, I31S, V115S, and I126S showed acceleration similar to that achieved with the Δ13-25 deletion and serine mutations within the hydrophobic patch of the PAS-cap α-helix (cf. **Figs. 4B** and **2C**). In contrast, I42S, V113S, and L123S, which exhibited faster compared to WT channels, were slower than the Δ13-25 deletion mutant. Within the CNBh domain, V795S, V798S and L825S mutants had fast deactivation like the Δ13-25 deletion mutant. The V_1/2_ of the mutant activation curves was not affected compared to WT (**Fig. S4** and **Table S5)**. These findings suggest that hydrophobic residues within the globular PAS and CNBh domains also contribute to the hydrophobic nexus sustaining slow deactivation. Based on their functional phenotypes, these hydrophobicity disrupting mutants can be divided into fast and intermediate deactivation groups.

**Fig. 4.**
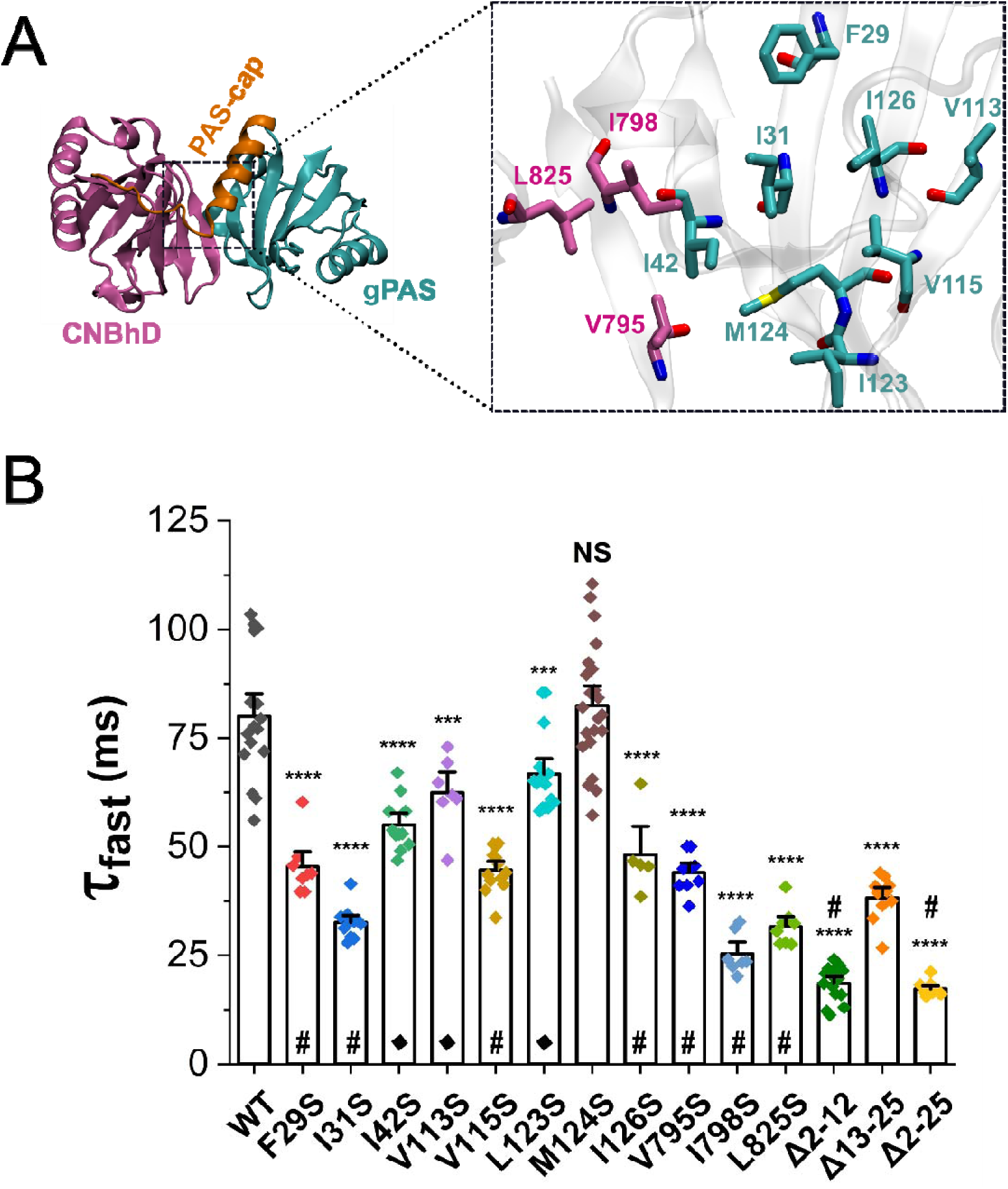
Residues constituting the hydrophobic nexus establish slow deactivation. (**A**) Representation of the PAS-CNBhD complex of hERG1a, highlighting the intracellular PAS-cap (orange), globular PAS (cyan), and CNBh (pink) domains. Inset shows hydrophobic residues at the globular PAS (F29, I31, I42, V113, V115, L123, M124, and I126) and CNBh (V795, I798, and L825) domains proposed to conform the three-dimensional nexus (PDB entry: 5VA1). (**B**) Fast components of deactivation in WT and F29S, I31S, I42S, V113S, V115S, L123S, M124S, I126S, V795S, I798S, L825S, Δ2-12, Δ13-25, and Δ2-25 mutant channels (****, P < 0.0001; ***, P < 0.001; NS: not significant, P > 0.05). Symbol “δ” indicates that the mutant deactivation rate is statistically faster compared to Δ13-25. Symbol “#” indicates that the mutant deactivation rate is not statistically significant compared to Δ13-25. Data are mean ± SEM.

### Perturbation of hydrophobicity predicts structural rearrangements at the nexus interface per MD simulations analysis

Our functional and structural analyses suggest that hydrophobic residues located at the PAS-cap α-helix, globular PAS and CNBh domains form a three-dimensional nexus that actively modulate hERG channel closing. To explore a possible mechanism by which disruption of nexus hydrophobicity yielded accelerated deactivation kinetics, we analyzed the nexus hydrophobic network throughout our fully atomistic MDS. We determined the average distance between the last carbon atom in the side chain of hydrophobic residues in the PAS-cap α-helix and their counterpart in proximal amino acids in the globular PAS and/or CNBh domains in the WT channel over the 175 ns MD simulation (**Fig. 5A, C, E**, and **G-I**). Similar analysis was performed for mutant channels using in this case the last carbon in the side chain of introduced serine residues (carbon β) (**Fig. 5B, D**, and **F**). Comparison of average distances between the slow (WT and F14S) and fast deactivating (L15S, I18S and I19S) channels was used as a proxy to determine whether structural changes within the hydrophobic nexus were induced by serine substitutions and related to the faster deactivation phenotype.

**Fig. 5.**
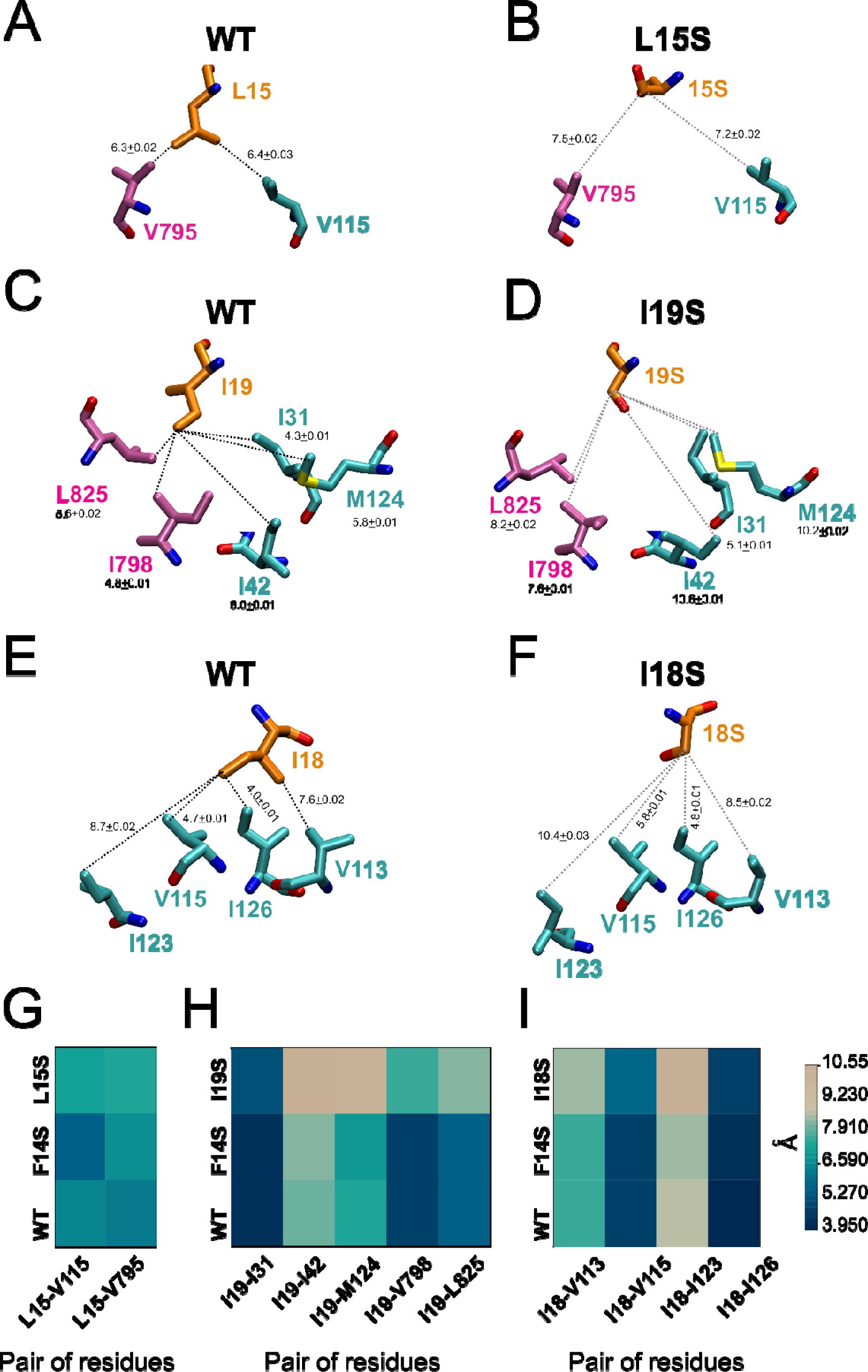
Serine substitutions of hydrophobic residues at the PAS-cap α-helix trigger structural changes within the hydrophobic nexus in hERG channels. Average distance between the residues L15 and I19 of the PAS-cap α-helix and residues at the globular PAS and CNBh domains in the WT (**A** and **C**, respectively), L15S (**B**), and I19S (**D**) channels. Mean distance between the residue in position 18 of the PAS-cap helix with residues at the globular PAS in the WT (**E**) and I18S (**F**) channels. Comparison of the average distance between hydrophobic residues at PAS-cap helix and amino acids at the globular PAS and/or CNBhD between the WT, F14S, and L15S (**G**), I19S (**H**) and I18S (**I**) channels (PDB entry: 5VA1).

Analysis *in silico* revealed that, when L15 is mutated to serine, average distances to residues located at the globular PAS (V115) and CNBh (V795) domains increase compared to slow deactivating WT and F14S channels, as indicated by color gradations corresponding to the heat map scale (**Fig. 5A-B, G**). I19S produced a similar effect, increasing the distances with residues I31, I42, and M124 at the globular PAS, and I798, and L825 at the CNBh domain (**Fig. 5C-D, H**). Finally, mutation of I18, which has close-contact residues only with the globular PAS domain, resulted in an increased distance to V113, V115, I123 and I126 (**Fig. 5E-F, I**). For a better visualization of structural rearrangements induced by serine mutations, **Fig. S5** shows surface representations of the PAS-cap α-helix, globular PAS and CNBh domains in orange, teal, and pink, respectively. In WT hERG, the PAS-cap α-helix, globular PAS, and CNBh domains form a tight central interface that constitutes the hydrophobic nexus (dashed white square in **Fig. S5A**). After introducing serine mutations at L15, I18, or I19, the hydrophobic nexus was compromised as suggested by the decreased surface contacts between the three domains in the central interface (dashed white square in **Fig. S5B- D**). Similar to our functional and α-helix secondary structure MDS analyses, the disruption of the nexus interface was more noticeable in I18S, as noted by decreased surface contact between the globular PAS and CNBh domains (black arrows in **Fig. S5D**).

We next evaluated if disrupting the nexus hydrophobicity produces rearrangements in the global positioning of the constituent domains. Here, we determined the average distance from the carbon α (C_α_) of hydrophobic residues at the globular PAS and CNBh domains to the C_α_ of residues 15, 18 and 19 in WT and mutant channels, as a proxy to backbone movements. Comparison of these values showed that serine substitutions do not cause significant changes in the relative position of the PAS-cap α-helix relative to the globular PAS and CNBh domains (**Fig. S6**).

These functional and MDS results suggest that the perturbation of PAS-cap α-helix hydrophobicity diminishes the probability of its interaction with proximate side chains of amino acids within the globular PAS and CNBh domains, triggering a faster current decay. Moreover, the accelerated deactivation appears to be due to discrete rearrangements of amino acid side chains at the nexus interface rather than macroscopic movements of the composing domains. Longer simulations may be required to capture such movements.

### Perturbation of the hydrophobic nexus results in movements of the PAS-cap tip away from the S4-S5 linker per MD simulation analysis

Gating of hERG channels has been proposed to occur via a “foot-in-the-door” mechanism, in which the close apposition between the PAS-cap N-terminus and the S4- S5 linker slows down the channel closing (Wang et al., 1998; Wang et al., 2000; de la Pena et al., 2011; Fernandez-Trillo et al., 2011; Ng et al., 2012; Hull et al., 2014; Wang and MacKinnon, 2017). Thus, if the PAS-cap unstructured segment serves as the “foot,” we reasoned that faster deactivation should be accompanied by its displacement away from the S4-S5 linker, its proposed site of interaction with the gating machinery (Wang et al., 1998; Wang et al., 2000; de la Pena et al., 2011). To test this hypothesis, we compared the proximity between C_α_ of residues V3 and D540, at the PAS-cap tip and S4-S5 linker respectively, in WT and fast deactivating mutants L15S, I18S, and I19S. The mutant channels showed an increased distance (approximately 2 Å) between the PAS-cap and S4-S5 linker when compared to WT throughout the simulation time (**Fig. 6**). These results suggest that changes within the hydrophobic nexus network caused by serine substitutions move the PAS-cap tip away from the S4-S5 linker, removing an allosteric barrier to channel closing.

**Fig. 6.**
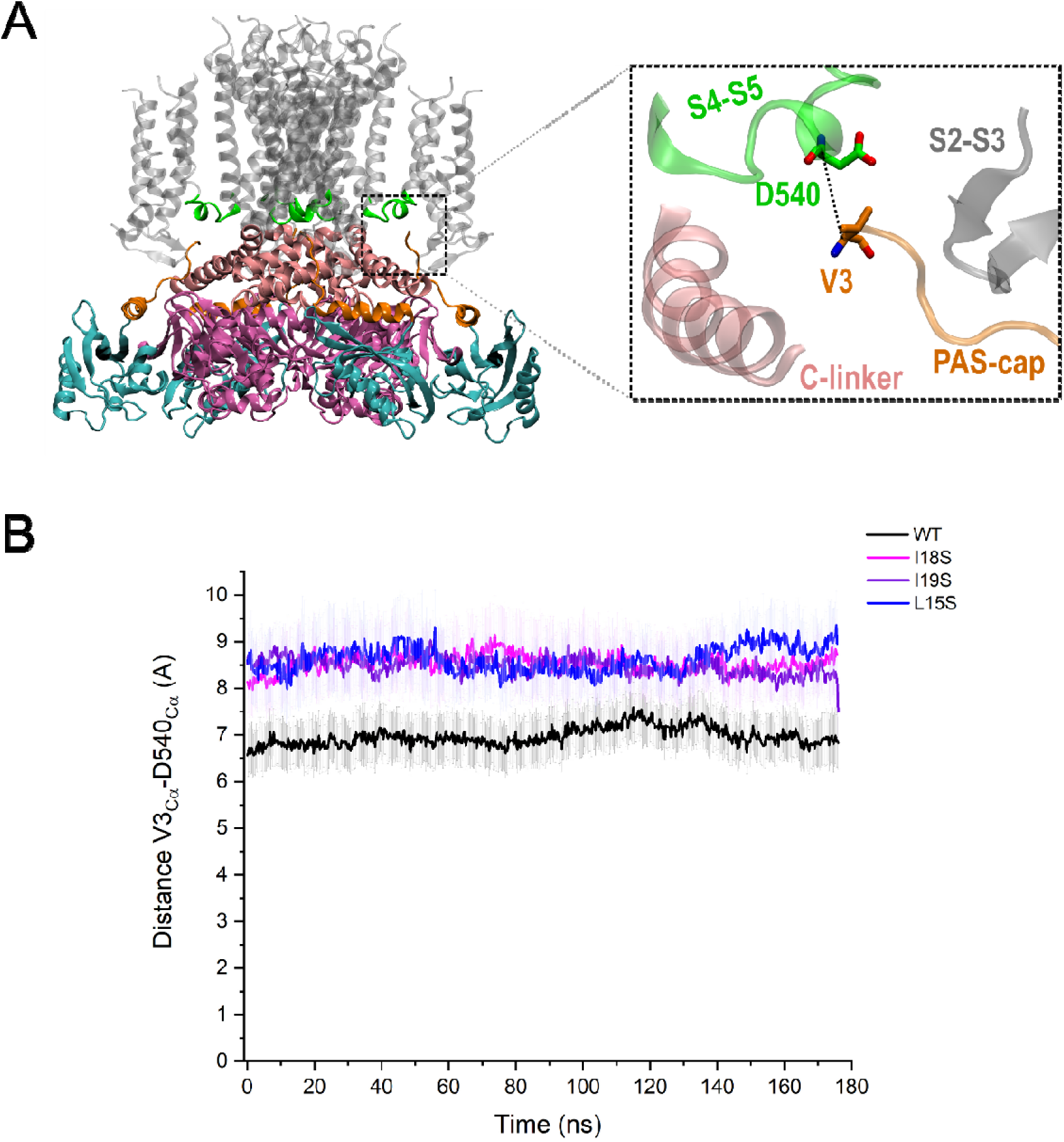
Perturbation of the nexus hydrophobicity results in decreased proximity between the PAS-cap tip and the S4-S5 linker. **(A)** Cryo-EM structure of a hERG1a tetramer (PDB entry: 5VA1) highlighting the intracellular PAS-cap (orange), globular PAS (gPAS) (cyan), the S4-S5 linker (green), C-linker (light pink), cyclic nucleotide-binding homology domain (CNBhD) (pink), and transmembrane domains (TMD) (grey) Inset shows amino acids used as a reference to determine the proximity between the PAS-cap tip (V3 in orange) and S4-S5 linker (D540 in green). The dashed line shows the measured distance. (**B**) Average distance between the Cα of residues V3 and D540 along the MDS.

## Discussion

In this study, we used structural and functional analyses, bioinformatics, mutagenesis, and MDS to define a hydrophobic nexus regulating slow deactivation in hERG channels. This region is formed by hydrophobic residues at the PAS-cap α-helix, the globular PAS, and CNBh domains. Perturbation of hydrophobicity of the constituent residues accelerated channel deactivation, largely without disrupting the PAS-cap helicity as determined by MDS. MDS also revealed that reduction of hydrophobicity (with serine substitutions) caused a rearrangement within the nexus leading to decreased contact at the interface between its constituent domains and increased distance between the PAS-cap tip and the S4-S5 linker, the proposed substrate for the “foot-in-the-door” that impedes channel closing. Based on our functional and MDS analyses, we propose that a hydrophobic network at the interface of the cytosolic PAS-cap α-helix, globular PAS, and CNBh domains is critical in establishing hERG channel slow deactivation. We speculate this hydrophobic nexus helps position the PAS-cap tip where it can stabilize the open state of the channel (Wang and MacKinnon, 2017; Robertson and Morais-Cabral, 2020).

The role of the PAS-cap in hERG channel gating has been studied using structural and functional approaches (Schonherr and Heinemann, 1996; Morais Cabral et al., 1998; Wang et al., 1998; Wang et al., 2000; Muskett et al., 2011; Ng et al., 2011; Tan et al., 2012; Ng et al., 2014), and several groups have reported important residue interactions between the tip of the PAS-cap unstructured region and the C-linker (Ng et al., 2014) and S4-S5 linker (Wang et al., 1998; de la Pena et al., 2011; Fernandez-Trillo et al., 2011; Ng et al., 2012; Hull et al., 2014; de la Pena et al., 2015). A role for the charged face of the PAS-cap α-helix, whose residues point to hydrophilic amino acids in the CNBh domain, was previously demonstrated, as channels with alanine or charge-reversal point mutations accelerated deactivation (Muskett et al., 2011; Ng et al., 2011). Alanine scanning of the PAS-cap (residues 2-25) produced mostly modest effects on deactivation and led to the conclusion that the PAS-cap α-helix acts as a physical spacer but not a key interactor with the globular PAS and CNBh domains (Ng et al., 2011). Given that serine substitutions phenocopy PAS-cap α-helix deletion, we propose that hydrophobic alanine substitutions in the previous study were insufficiently disruptive of the essentially hydrophobic nature of the complex.

Structural differences at the hydrophobic nexus may explain the dramatic disparities in deactivation kinetics between ERG and the related EAG1 channels (cf. **Fig. 1**).

Comparison of the sequence of the motifs constituting the tripartite nexus in mammalian ERG and EAG1 indicates reduced hydrophobicity comporting with the faster deactivation in EAG1. This prediction is corroborated by the cryo-EM structures showing reduced hydrophobicity and a more open conformation of the corresponding tripartite region in rEAG1 compared to hERG (Whicher and MacKinnon, 2016; Wang and MacKinnon, 2017). In support of this assumption, Ng et al. showed that neither the PAS-cap nor the globular PAS domains of EAG1 channels are functionally interchangeable with the respective N-terminal domains in hERG channels (Ng et al., 2014). Introducing the hERG PAS-cap, globular PAS, C-linker, and CNBh domains (i.e., the hERG hydrophobic nexus) into an EAG1-hERG chimera, however, results in slowed deactivation kinetics characteristic of hERG (Whicher and MacKinnon, 2019). These studies confirm the hERG PAS-cap is required to slow hERG channel closing and suggest that incomplete reconstitution of the three-dimensional hydrophobic nexus is insufficient to slow closing kinetics.

Comparison of fast deactivation of PAS-cap truncated hERG channels (i.e., Δ2-12, Δ13- 25, and Δ2-25) with the serine substitutions at either the PAS-cap α-helix, globular PAS, or CNBh domains showed that a majority of the point mutations phenocopied the rate of acceleration resulting from the PAS-cap α-helix deletion (Δ13-25), rather than the maximal acceleration shown by deletion of the PAS-cap disordered region (Δ2-12) or the entire PAS-cap (Δ2-25) (cf. **Fig. 2** and **Fig. 4**). This agrees with previous findings showing that residues at the tip of the PAS-cap disordered region are key mediators of hERG deactivation via specific interactions with the C-linker and the S4-S5 linker, where they may act as a “foot-in-the-door” to stabilize the open channel state (Wang et al., 1998; Wang et al., 2000; Fernandez-Trillo et al., 2011; Muskett et al., 2011; Ng et al., 2011; Ng et al., 2014; de la Pena et al., 2015). The intermediate deactivation phenotype conferred by hydrophobic nexus mutations leads us to conclude that the extreme N-terminal residues can interact with the channel gating machinery but do so with less efficacy if the hydrophobic nexus is disrupted. Thus, the identified three-dimensional nexus acts as a functional modulatory unit conferring slow deactivation within the allosteric pathway underlying hERG channel gating.

MDS analysis of the relative positioning of the side chain of hydrophobic residues within the nexus in WT and fast-deactivating hERG mutants showed that serine point mutations dissociate the tripartite region (cf. **Fig. 5** and **Fig. S5**) and trigger a faster current decay. Interestingly, residues I42, V113, and L123, whose serine mutations had the lowest effect on deactivation kinetics, showed the highest distance relative to hydrophobic residues at the PAS-cap α-helix (cf. **Fig. 4** and **Fig. 5**), suggesting that the closer the residue to the center of the hydrophobic nexus the more critical is its modulatory role. We reason that the integrity of the hydrophobic nexus serves to ensure the correct positioning of the PAS-cap domain to modulate gating. In support of this interpretation, recent work by Harley et al. showed that the globular PAS-CNBh domain interface is dynamic, and structural alterations at this interface affect the equilibrium between the open and closed states of the channel (Harley et al., 2021). Thus, decreased contacts between the domains forming the hERG hydrophobic nexus after introducing mutations at this site is consistent with the proposed state-dependent interactions between the PAS and CNBh domains. Our MDS analyses also suggest that discrete changes within the hydrophobic nexus network can trigger substantial distancing between the PAS-cap disordered region and the S4-S5 linker. Interestingly, this occurred even in the absence of macroscopic changes in the position of the constituent domains. Future studies including other domains possibly involved in the allosteric pathway (i.e., N-linker, S2-S3 linker, and C-linker) must be addressed to answer how these microscopic changes are transmitted to the PAS-cap tip.

The identified hydrophobic nexus not only sheds light onto the mechanism of hERG deactivation, but also provides insights into the mechanism of LQTS clinically observed in patients harboring hERG mutations. Several LQTS-associated point mutations have been identified at all components of the hydrophobic nexus (Haitin et al., 2013), with some affecting hERG function, domain stability, and/or channel trafficking (Anderson et al., 2006; Foo et al., 2019). LQTS-associated mutations within the hydrophobic patch of the globular PAS include F29L, I31S, and I42N, all of which are trafficking deficient (Anderson et al., 2006; Ke et al., 2013; Foo et al., 2019). Surface hydrophobic interactions are important during protein folding and assembly and are crucial in the stabilization of proteins (Pace et al., 2011). Thus, proper assembly of the hydrophobic nexus may be required to pass quality-control checkpoints in the endoplasmic reticulum (Zarei et al., 2004), possibly reflecting its importance in the fitness of the organism.

Globular PAS domain LQTS-associated mutants F29L, I42N, and M124R also hasten deactivation (Morais Cabral et al., 1998; Chen et al., 1999; Shushi et al., 2005; Ke et al., 2013). The hydrophobic nexus is therefore crucial for proper hERG channel trafficking and function, as several disease-associated mutations are localized within this modulatory unit.

In cardiomyocytes, where native I_Kr_ channels are thought to be heteromers of hERG1a and 1b isoforms (Jones et al., 2004; Sale et al., 2008; Jones et al., 2014), deactivation is faster than in hERG1a homomers but sufficiently slow to allow the recovery from inactivation and the resurgence of the tail current during repolarization (Sale et al., 2008). There will be fewer than four hydrophobic nexuses owing to the absence of PAS domains in hERG1b (London et al., 1997). Although the stoichiometry of native channels is not yet known, their relatively fast kinetics are consistent with fewer functional PAS domains (London et al., 1997; Trudeau et al., 2011; Gustina and Trudeau, 2013); conversion to 1a-like currents can be achieved by expressing the hERG1a PAS domain as a fragment in trans in cardiomyocytes (Jones et al., 2014). The relative contributions of the hydrophobic nexus in heteromeric hERG1a/1b channels remains to be determined.

In summary, our study proposes the three-dimensional hydrophobic nexus as a critical molecular determinant of hERG slow deactivation. Hydrophobic interactions between the PAS-cap, globular PAS and CNBh domains maintain the inter-subunit intracellular gating ring in a conformational arrangement where the PAS-cap is primed for the stabilization of the open state. Whether the PAS-cap dissociates from the hydrophobic nexus during gating remains to be demonstrated experimentally. We conclude that the PAS-cap α-helix has an active role mediating slow deactivation of hERG channels and that the three-dimensional hydrophobic nexus serves an important biophysical and physiological role in hERG gating.

## Materials and methods

### Molecular biology

The human ERG1a (hERG or KCNH2, GenBank accession no. <u>Q12809</u>) cDNA was cloned into the pGH19 oocyte expression vector. Mutations in WT hERG1a cDNA were made using the QuikChange site-directed mutagenesis kit (Agilent Technologies) and verified by DNA sequence analysis. WT and mutant DNA plasmids were linearized by HindIII-HF or EcoRI-HF and purified with the QIAquick PCR Purification Kit (QIAGEN). cRNA was transcribed *in vitro* with the mMESSAGE mMACHINE T7 kit (Thermo Fisher Scientific), purified, and solubilized with RNase-free water.

### Oocyte isolation and cRNA injection

Oocytes from *Xenopus laevis* female frogs [obtained from Nasco (Fort Atkinson, WI) and Xenopus1 Corp. (Dexter, MI)] were surgically removed, isolated manually, and treated with 75 μg/mL Liberase TM research-grade enzyme (Sigma-Aldrich) for 40–60 min in Ca^2+^-free ND96 solution [96 mM NaCl, 2 mM KCl, 1 mM MgCl_2_, and 5 mM HEPES, with pH adjusted to 7.4 with NaOH]. Defolliculated Stage IV and V oocytes were injected with WT or mutant cRNA and incubated for 1–5 d at 16°C in ND96 storage solution [96 mM NaCl, 2 mM KCl, 1 mM CaCl_2_, 1 mM MgCl_2_, 5 mM HEPES, 100mg/L bovine serum albumin, and 10 mg/L gentamycin, with pH adjusted to 7.4 with NaOH].

### Two-electrode voltage clamp

Oocytes expressing channels of interest were subjected to a standard two-microelectrode voltage clamp. All recordings were performed at room temperature (21°C ± 2 °C) and held at -80 mV between recordings. Leak subtraction was done offline using Ohm’s Law based on passive leak current measured at -80 mV. The resistance of the electrodes ranged from 0.5 to 2.0 MΩ in 2 M KCl. Recording solution [5 mM KCl, 93 mM N-methyl-D-glucamine, 1 mM MgCl_2_, 0.3 mM CaCl_2_, and 5 mM HEPES, with pH adjusted to 7.4 with MES] was perfused through the bath chamber during the recording.

Deactivation tail currents were evoked at –120 mV after a 4-s depolarizing step to +60 mV. The deactivating tail current was fit to a second order exponential Chebyshev fit equation 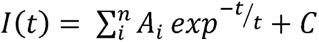 to obtain the fast time constant (τ) of deactivation.

Isochronal 1-s fitting measurements were obtained from the peak of the tail current, with the cursor of the fitting window advanced to the first point in time that did not force a fit to the recovery phase and extrapolated back to the moment of the voltage change. The deactivation traces were best fit with two ’s. At more negative voltages, most of the current was described by a single, fast deactivation tau, such that τ*_fast_* ≅ 1⁄*α* + *β*, corresponding to the sum of activation and deactivation, respectively. Since activation is negligible at negative voltages, τ*_fast_* ≅ 1/*β*. Thus, the reported time constants correspond to the dominant fast component, which contributes ∼90% of the deactivating current. Steady-state activation curves were generated using a pulse protocol with 1-s voltage steps ranging from –100 to +70 mV in 10 mV increments. Tail currents were measured during a subsequent step to –105 mV for 3 s. Normalized conductance values (I/I_max_) were obtained by normalizing the tail currents by the maximum tail current at +70 mV and were plotted as a function of the preceding voltage step. The resulting relation was fit with a Boltzmann equation *I*(*t*)=[(*V*-*V_rev_*)\**I_max_*]/(1+*exp*^(*V*-*V*_1/2_/*k*)^ to obtain the steady-state voltage dependence of activation parameters. Data were analyzed using pCLAMP 10.4 software (Molecular Devices), Origin 2021b (OriginLab) and GraphPad Prism 9.0 Version for Windows (GraphPad Software, Inc., CA, US). Results were considered statistically significant when p < 0.05. No blinding or randomization was conducted for the experiments. The Grubbs’ test was used to detect and exclude outliers in the data (GraphPad Software, Inc., CA, US). Data are shown as mean ± SE.

### Sequence data and analysis

We retrieved 20 sequences of ERG1a and 20 sequences of the long isoform of EAG1 channel in mammalian representative species. Protein sequences were obtained from the NCBI database (refseq_genomes, htgs, and wgs) using tbalstn with default settings. Protein sequences were aligned using the FFT-NS-1strategy from MAFFT v.745. Percent of identity for amino acids forming the interface between the PAS-cap, globular PAS and CNBh domains was calculated in Jalview (Waterhouse et al., 2009). The percentage of hydrophobic residues was determined using the Kyte-Doolittle scale available in Jalview. Residues facing the tripartite nexus were identified via the available structural models for tetrameric hERG1a (PDB:5VA1) and rEAG1 (PDB: 5K7L). Percent hydrophobicity was calculated as: 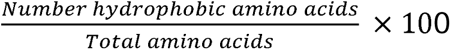 for ERG1 and EAG1 channels. Jalview was also used to generate multiple sequence alignment images.

### Molecular dynamics simulations

The three-dimensional structure of human ERG1 (PDB: 5VA1) was used for generating *in silico* serine substitutions at position F14, L15, I18 and I19 by using the plugin Residue and loop mutation of the Maestro-Schrödinger suite (2018-4 Schrödinger Release 2018-2: Maestro, Schrödinger, LLC, New York, NY,2018). The WT and mutant channels were prepared at pH 7.0 and subjected to energy minimization in vacuum through the Maestro-Schrödinger suite (Schrödinger Release 2018-2: Maestro, Schrödinger, LLC, New York, NY,2018). The channels were embedded into a pre-equilibrated POPC lipid bilayer in an orthorhombic box with periodic borders, filled with Single Point Charge (SPC) water molecules and an ionic concentration of 150 mM KCl. Full atom molecular dynamics simulations of 300ns were performed, per triplicates, in a NPT ensemble (P = 1 atm, T =310 K) with the Desmond software using the OPLS v.2005 force field (40). Structures were collected every 0.2 ns during the MDS having 1113 frames per simulation. Root mean squared deviation (RMSD) of the protein backbone was calculated using RMSD trajectory tool of VMD 1.9.3 and used to determine the time when the system reaches equilibrium. The simulation equilibrium time was achieved after 125 ns in all conditions; therefore, the last 175 ns were used for data analysis. Analyses of the secondary structure were performed using timeline plugin in VMD 1.9.3 (Humphrey et al., 1996) at equilibrium. Mean interaction distance between the residues was determined by averaging the measured distance between the last carbon atom of residues in close contact along the simulation time in VMD 1.9.3. The same approach was used to determine the distance between alpha carbons. VMD1.9.3 was also used to generate the figures.

## Acknowledgements

Work was supported by NIH grants 1R01NS081320 (GAR) and F99NS12582 (WASS). This research was supported by the National Institutes of Health, under Ruth L. Kirschstein National Research Service Award T32 HL 007936 from the National Heart Lung and Blood Institute to the University of Wisconsin-Madison Cardiovascular Research Center (LFA). DB thanks to Fondecyt de Iniciación a la Investigación N° 11220444. The authors thank Dr. Joao Morais-Cabral for helpful discussion.

## Author contributions

Conception and design of the work: L.F.-A, W.A.S.-S., and G.A.R. Acquisition of data and analysis: W.A.S.-S., L.F-A, D.B., and J.L. Interpretation of data: L.F.-A., W.A.S.-S., and G.A.R. Draft and/or revision of manuscript and final approval: W.A.S.-S., L.F.-A., D.B., L.D., and G.A.R.

