## Supplementary Materials for "An intracellular hydrophobic nexus is critical for slow deactivation in hERG channels"

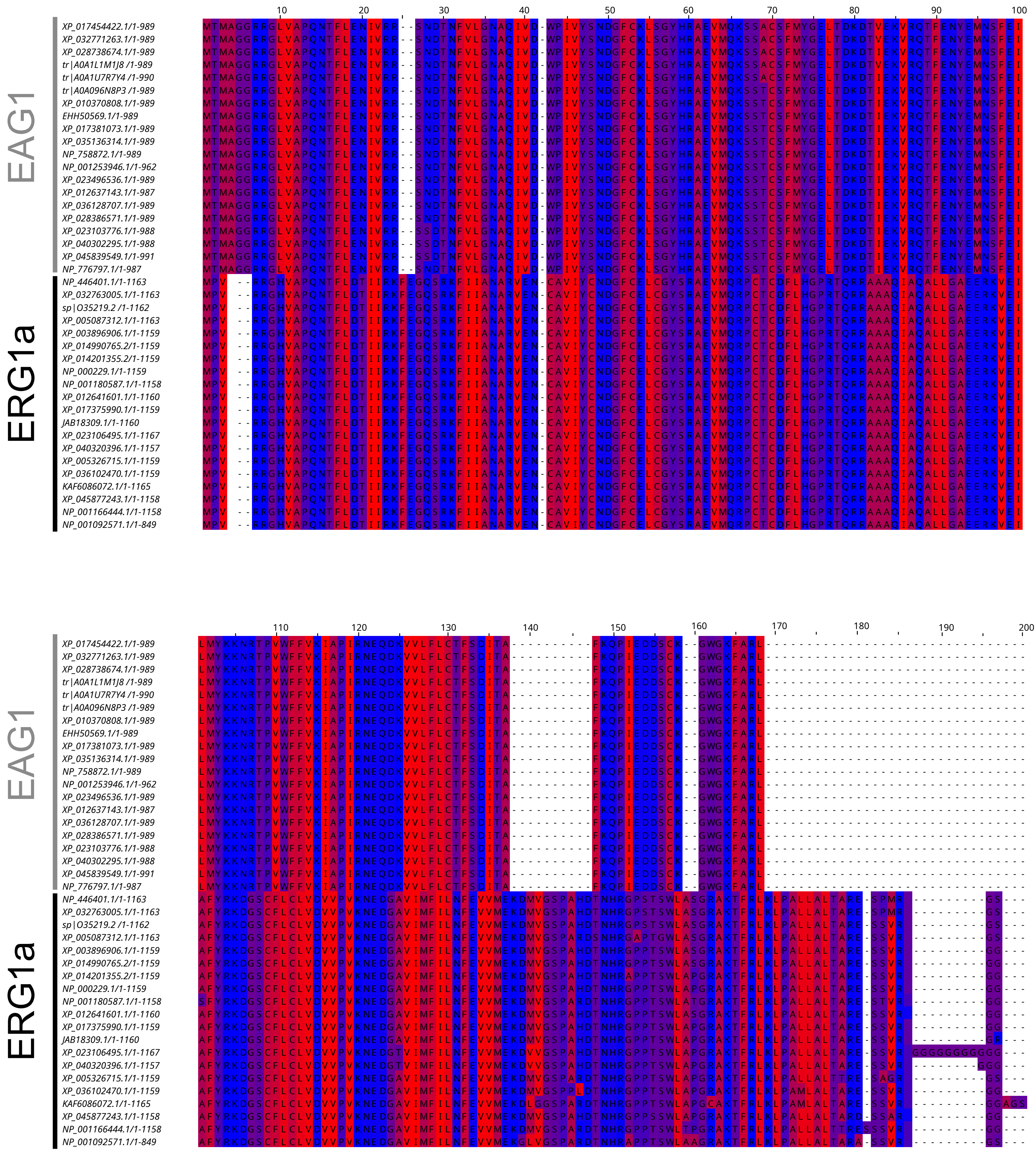
EAG1
ERG1a
10
20
30
40
50
60
70
80
90
100
110
120
130
140
150
160
170
180
190
200
XP_017454422.1/1-989
XP_032771263.1/1-989
XP_028738674.1/1-989
tr|A0A1L1M1J8/1-989
tr|A0A1U7R7Y4/1-990
tr|A0A096N8P3/1-989
XP_010370808.1/1-989
EHH50569.1/1-989
XP_017381073.1/1-989
XP_035136314.1/1-989
NP_758872.1/1-989
NP_001253946.1/1-962
XP_023496536.1/1-989
XP_012637143.1/1-987
XP_036128707.1/1-989
XP_028386571.1/1-989
XP_023103776.1/1-988
XP_040302295.1/1-988
XP_045839549.1/1-991
NP_776797.1/1-987
NP_446401.1/1-1163
XP_032763005.1/1-1163
sp|O35219.2/1-1162
XP_005087312.1/1-1163
XP_003896906.1/1-1159
XP_014990765.2/1-1159
XP_014201355.2/1-1159
NP_000229.1/1-1159
NP_001180587.1/1-1158
XP_012641601.1/1-1160
XP_017375990.1/1-1159
JAB18309.1/1-1160
XP_023106495.1/1-1167
XP_040320396.1/1-1157
XP_005326715.1/1-1159
XP_036102470.1/1-1159
KAF6086072.1/1-1165
XP_045877243.1/1-1158
NP_001166444.1/1-1158
NP_001092571.1/1-849

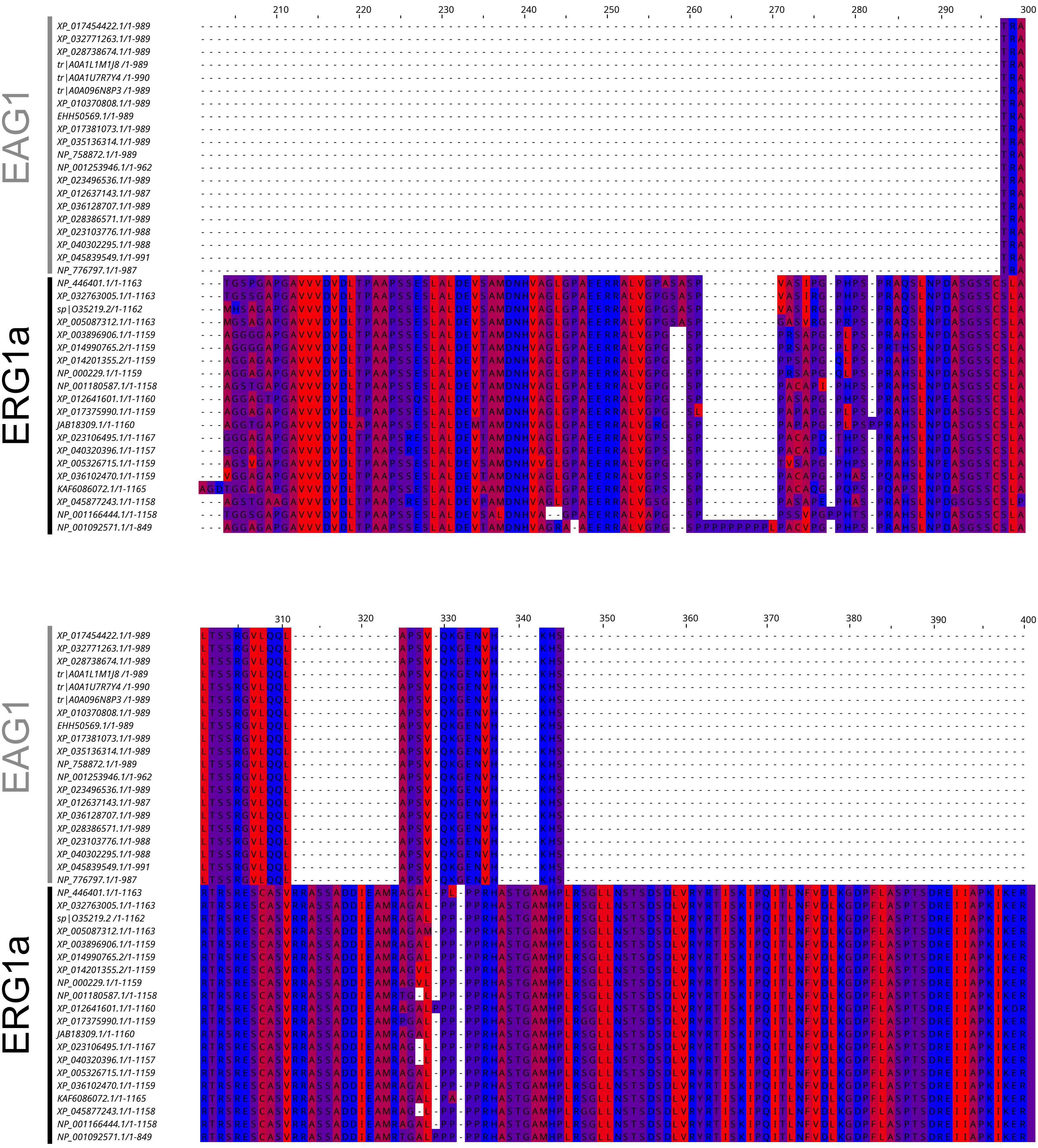

210
220
230
240
250
260
270
280
290
300
EAG1
XP_017454422.1/1-989
XP_032771263.1/1-989
XP_028738674.1/1-989
tr|A0A1L1M1J8 /1-989
tr|A0A1U7R7Y4 /1-990
tr|A0A096N8P3 /1-989
XP_010370808.1/1-989
EHH50569.1/1-989
XP_017381073.1/1-989
XP_035136314.1/1-989
NP_758872.1/1-989
NP_001253946.1/1-962
XP_023496536.1/1-989
XP_012637143.1/1-987
XP_036128707.1/1-989
XP_028386571.1/1-989
XP_023103776.1/1-988
XP_040302295.1/1-988
XP_045839549.1/1-991
NP_776797.1/1-987
ERG1a
NP_446401.1/1-1163
XP_032763005.1/1-1163
sp|O35219.2/1-1162
XP_005087312.1/1-1163
XP_003896906.1/1-1159
XP_014990765.2/1-1159
XP_014201355.2/1-1159
NP_000229.1/1-1159
NP_001180587.1/1-1158
XP_012641601.1/1-1160
XP_017375990.1/1-1159
JAB18309.1/1-1160
XP_023106495.1/1-1167
XP_040320396.1/1-1157
XP_005326715.1/1-1159
XP_036102470.1/1-1159
KAF6086072.1/1-1165
XP_045877243.1/1-1158
NP_001166444.1/1-1158
NP_001092571.1/1-849
310
320
330
340
350
360
370
380
390
400
EAG1
ERG1a

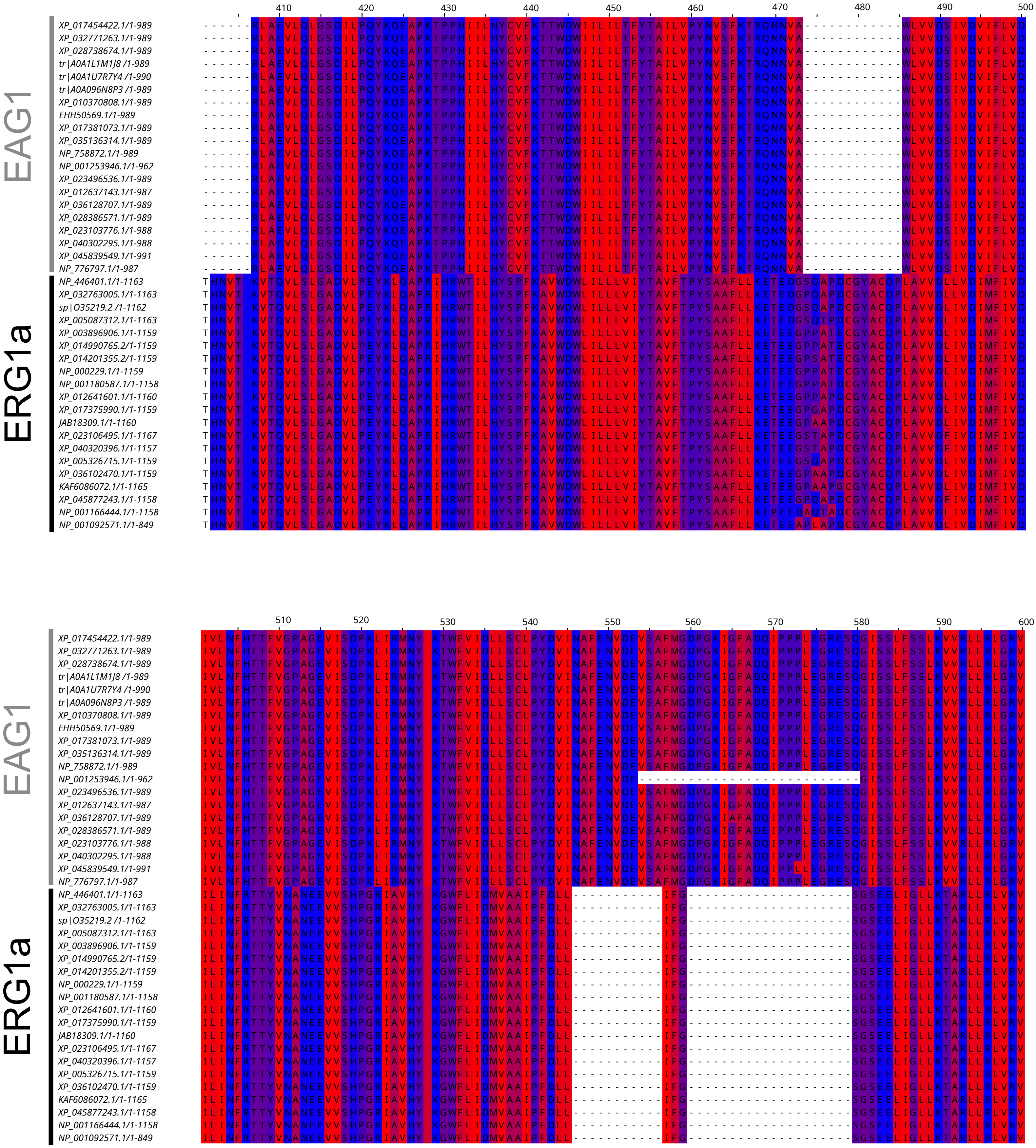

EAG1
ERG1a
410
420
430
440
450
460
470
480
490
500
510
520
530
540
550
560
570
580
590
600
XP_017454422.1/1-989
XP_032771263.1/1-989
XP_028738674.1/1-989
tr|A0A1L1M1J8 /1-989
tr|A0A1U7R7Y4 /1-990
tr|A0A096N8P3 /1-989
XP_010370808.1/1-989
EHH50569.1/1-989
XP_017381073.1/1-989
XP_035136314.1/1-989
NP_758872.1/1-989
NP_001253946.1/1-962
XP_023496536.1/1-989
XP_012637143.1/1-987
XP_036128707.1/1-989
XP_028386571.1/1-989
XP_023103776.1/1-988
XP_040302295.1/1-988
XP_045839549.1/1-991
NP_776797.1/1-987
NP_446401.1/1-1163
XP_032763005.1/1-1163
sp|O35219.2 /1-1162
XP_005087312.1/1-1163
XP_003896906.1/1-1159
XP_014990765.2/1-1159
XP_014201355.2/1-1159
NP_000229.1/1-1159
NP_001180587.1/1-1158
XP_012641601.1/1-1160
XP_017375990.1/1-1159
JAB18309.1/1-1160
XP_023106495.1/1-1167
XP_040320396.1/1-1157
XP_005326715.1/1-1159
XP_036102470.1/1-1159
KAF6086072.1/1-1165
XP_045877243.1/1-1158
NP_001166444.1/1-1158
NP_001092571.1/1-849

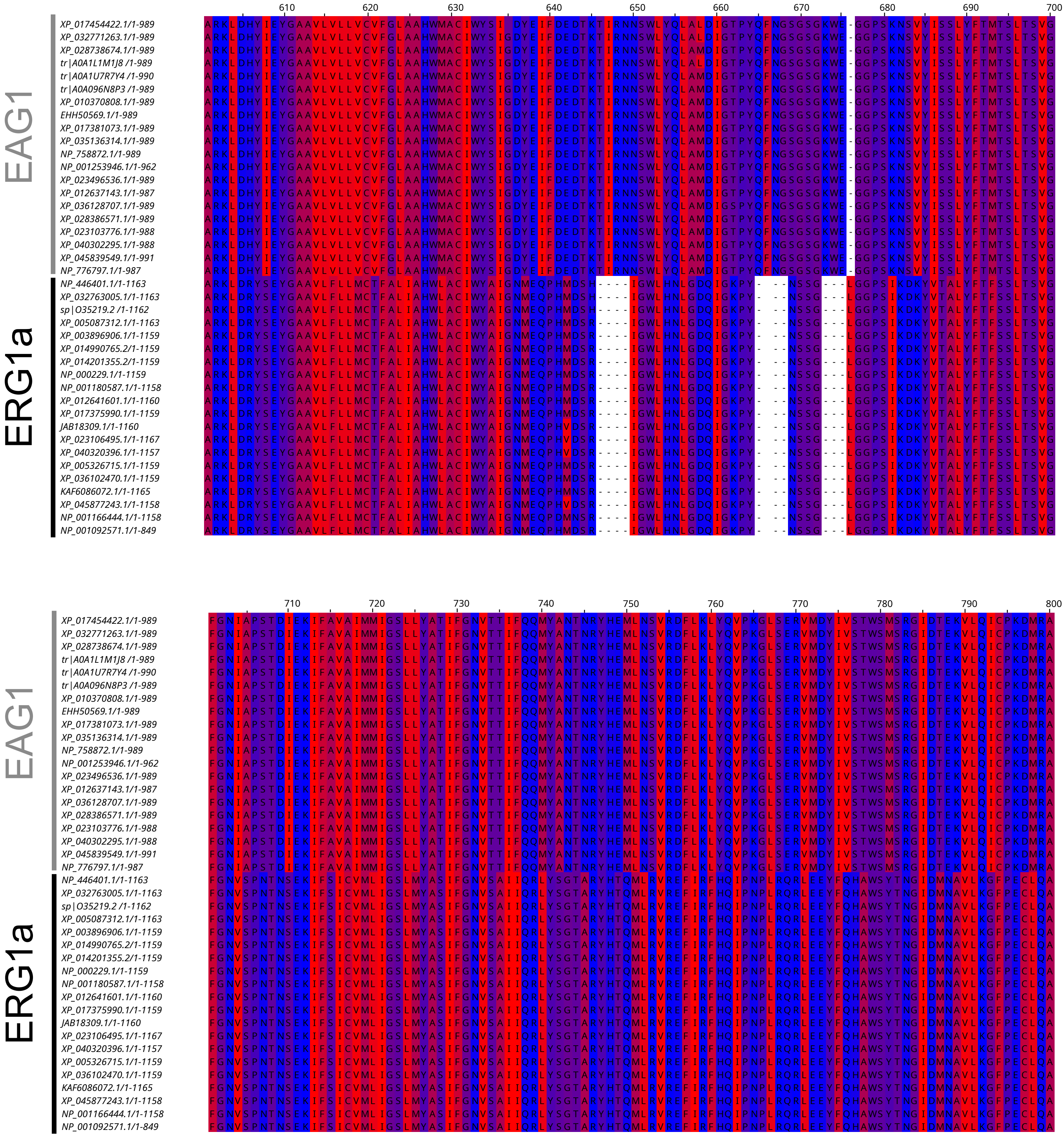
EAG1
ERG1a
610
620
630
640
650
660
670
680
690
700
710
720
730
740
750
760
770
780
790
800
XP_017454422.1/1-989
XP_032771263.1/1-989
XP_028738674.1/1-989
tr|A0A1L1M1J8 /1-989
tr|A0A1U7R7Y4 /1-990
tr|A0A096N8P3 /1-989
XP_010370808.1/1-989
EHH50569.1/1-989
XP_017381073.1/1-989
XP_035136314.1/1-989
NP_758872.1/1-989
NP_001253946.1/1-962
XP_023496536.1/1-989
XP_012637143.1/1-987
XP_036128707.1/1-989
XP_028386571.1/1-989
XP_023103776.1/1-988
XP_040302295.1/1-988
XP_045839549.1/1-991
NP_776797.1/1-987
NP_446401.1/1-1163
XP_032763005.1/1-1163
sp|O35219.2 /1-1162
XP_005087312.1/1-1163
XP_003896906.1/1-1159
XP_014990765.2/1-1159
XP_014201355.2/1-1159
NP_000229.1/1-1159
NP_001180587.1/1-1158
XP_012641601.1/1-1160
XP_017375990.1/1-1159
JAB18309.1/1-1160
XP_023106495.1/1-1167
XP_040320396.1/1-1157
XP_005326715.1/1-1159
XP_036102470.1/1-1159
KAF6086072.1/1-1165
XP_045877243.1/1-1158
NP_001166444.1/1-1158
NP_001092571.1/1-849

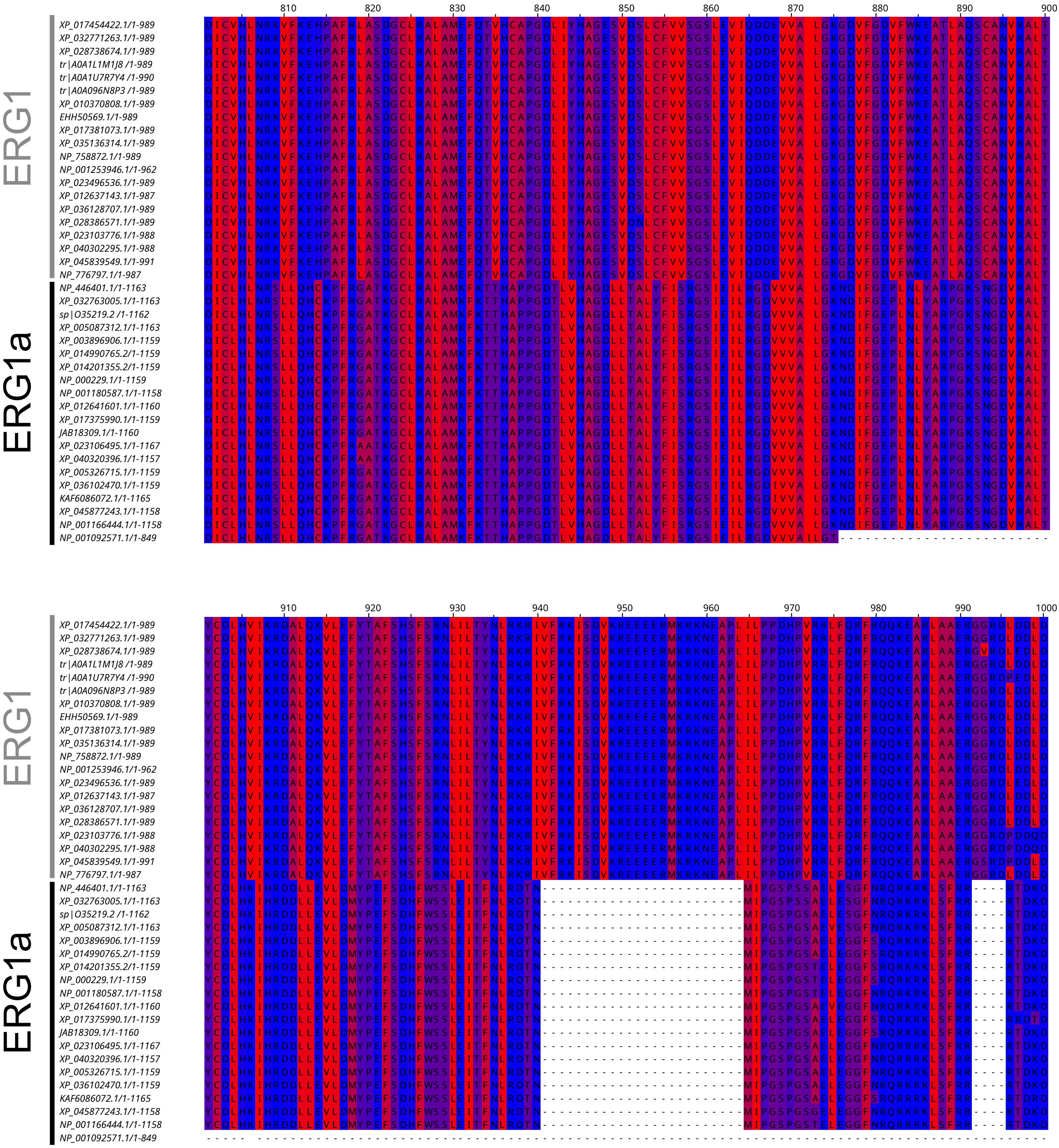

810
820
830
840
850
860
870
880
890
900
ERG1
ERG1a
XP_017454422.1/1-989
XP_032771263.1/1-989
XP_028738674.1/1-989
tr|A0A1L1M1J8 /1-989
tr|A0A1U7R7Y4 /1-990
tr|A0A096N8P3 /1-989
XP_010370808.1/1-989
EHH50569.1/1-989
XP_017381073.1/1-989
XP_035136314.1/1-989
NP_758872.1/1-989
NP_001253946.1/1-962
XP_023496536.1/1-989
XP_012637143.1/1-987
XP_036128707.1/1-989
XP_028386571.1/1-989
XP_023103776.1/1-988
XP_040302295.1/1-988
XP_045839549.1/1-991
NP_776797.1/1-987
NP_446401.1/1-1163
XP_032763005.1/1-1163
sp|O35219.2 /1-1162
XP_005087312.1/1-1163
XP_003896906.1/1-1159
XP_014990765.2/1-1159
XP_014201355.2/1-1159
NP_000229.1/1-1159
NP_001180587.1/1-1158
XP_012641601.1/1-1160
XP_017375990.1/1-1159
JAB18309.1/1-1160
XP_023106495.1/1-1167
XP_040320396.1/1-1157
XP_005326715.1/1-1159
XP_036102470.1/1-1159
KAF6086072.1/1-1165
XP_045877243.1/1-1158
NP_001166444.1/1-1158
NP_001092571.1/1-849
910
920
930
940
950
960
970
980
990
1000

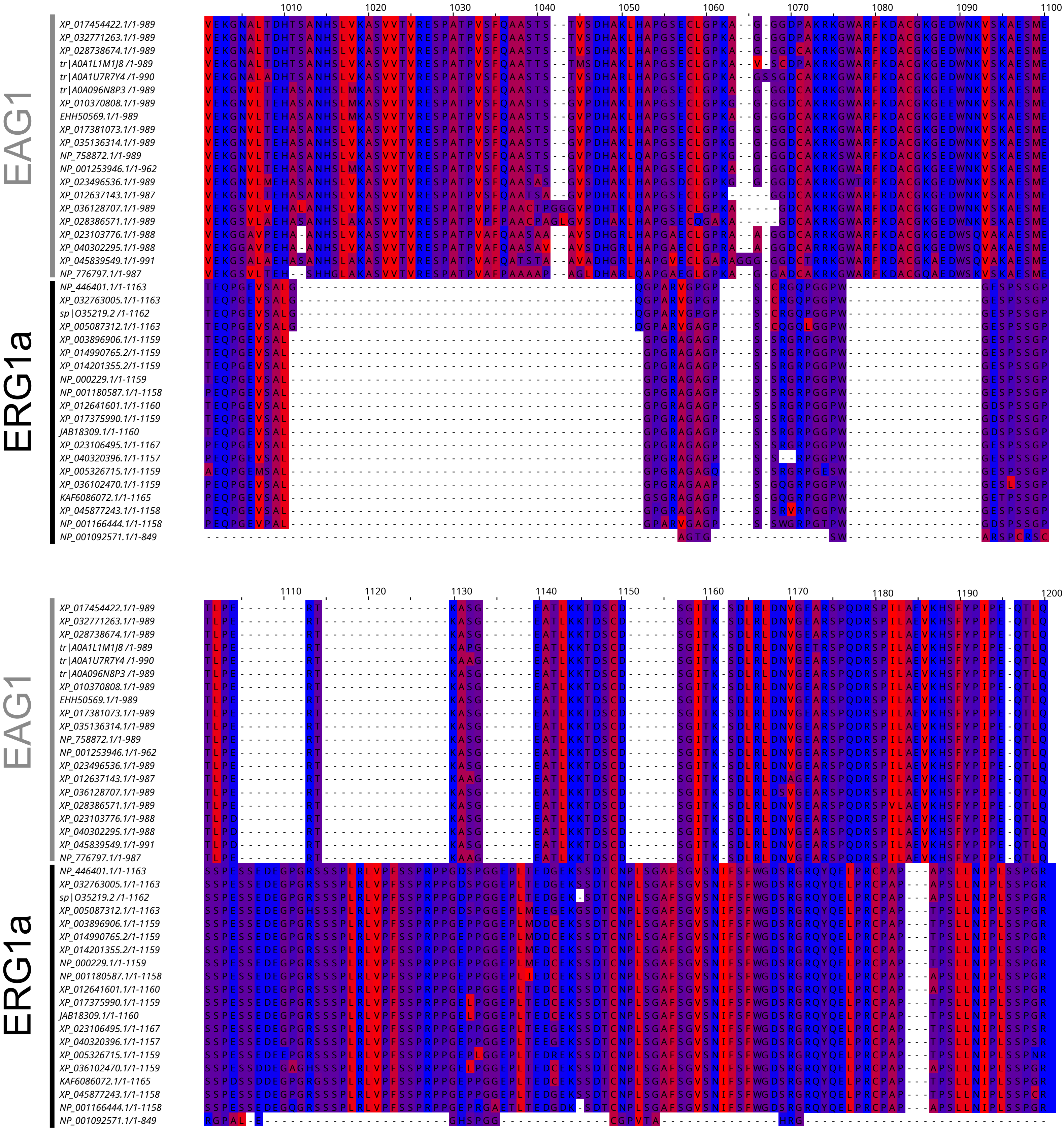
1010
1020
1030
1040
1050
1060
1070
1080
1090
1100
EAG1
ERG1a
XP_017454422.1/1-989
XP_032771263.1/1-989
XP_028738674.1/1-989
tr|A0A1L1M1J8/1-989
tr|A0A1U7R7Y4/1-990
tr|A0A096N8P3/1-989
XP_010370808.1/1-989
EHH50569.1/1-989
XP_017381073.1/1-989
XP_035136314.1/1-989
NP_758872.1/1-989
NP_001253946.1/1-962
XP_023496536.1/1-989
XP_012637143.1/1-987
XP_036128707.1/1-989
XP_028386571.1/1-989
XP_023103776.1/1-988
XP_040302295.1/1-988
XP_045839549.1/1-991
NP_776797.1/1-987
NP_446401.1/1-1163
XP_032763005.1/1-1163
sp|O35219.2/1-1162
XP_005087312.1/1-1163
XP_003896906.1/1-1159
XP_014990765.2/1-1159
XP_014201355.2/1-1159
NP_000229.1/1-1159
NP_001180587.1/1-1158
XP_012641601.1/1-1160
XP_017375990.1/1-1159
JAB18309.1/1-1160
XP_023106495.1/1-1167
XP_040320396.1/1-1157
XP_005326715.1/1-1159
XP_036102470.1/1-1159
KAF6086072.1/1-1165
XP_045877243.1/1-1158
NP_001166444.1/1-1158
NP_001092571.1/1-849
1110
1120
1130
1140
1150
1160
1170
1180
1190
1200

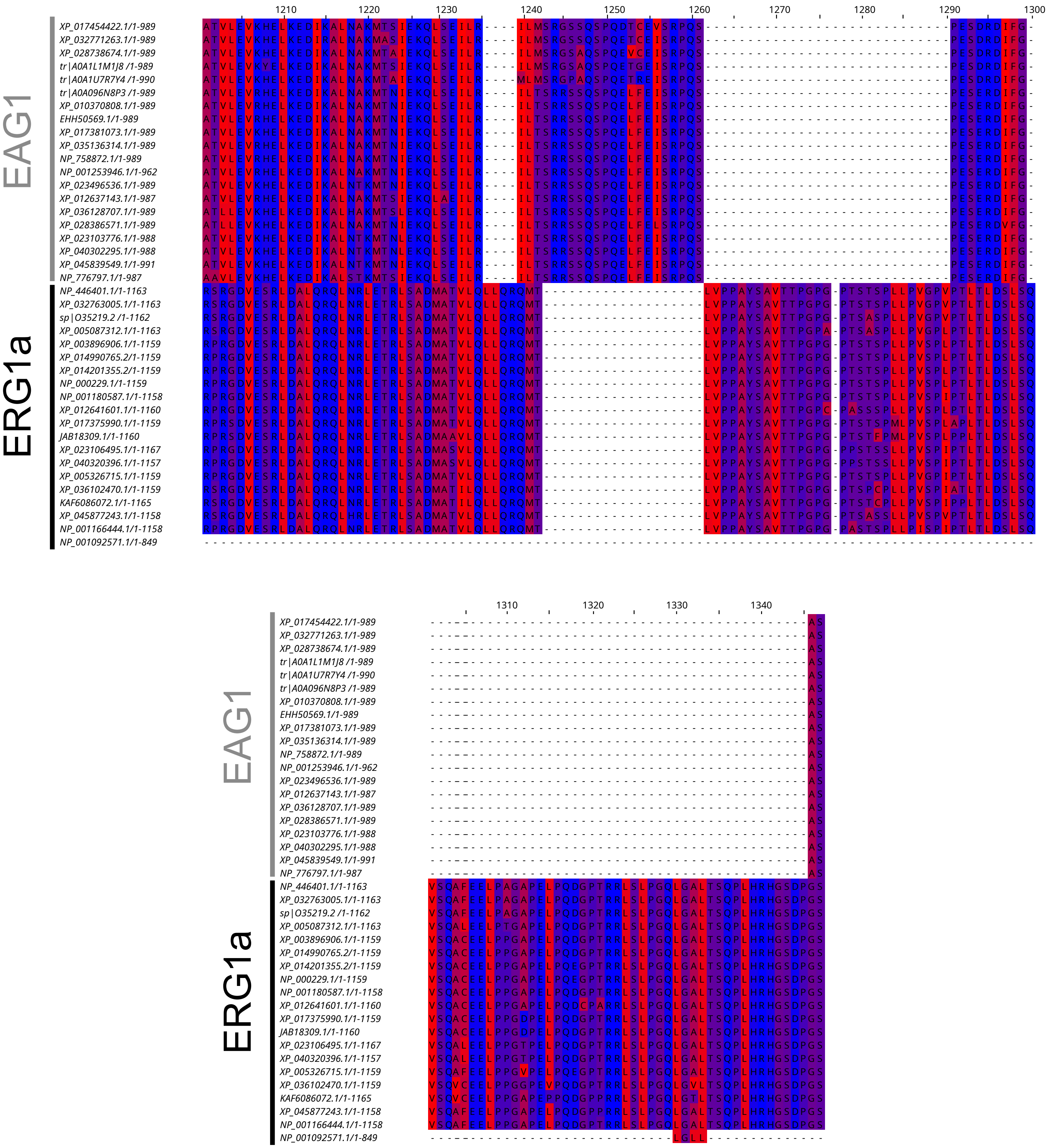
1210
1220
1230
1240
1250
1260
1270
1280
1290
1300
EAG1
XP_017454422.1/1-989
XP_032771263.1/1-989
XP_028738674.1/1-989
tr|A0A1L1M1J8 /1-989
tr|A0A1U7R7Y4 /1-990
tr|A0A096N8P3 /1-989
XP_010370808.1/1-989
EHH50569.1/1-989
XP_017381073.1/1-989
XP_035136314.1/1-989
NP_758872.1/1-989
NP_001253946.1/1-962
XP_023496536.1/1-989
XP_012637143.1/1-987
XP_036128707.1/1-989
XP_028386571.1/1-989
XP_023103776.1/1-988
XP_040302295.1/1-988
XP_045839549.1/1-991
NP_776797.1/1-987
ERG1a
NP_446401.1/1-1163
XP_032763005.1/1-1163
sp|O35219.2 /1-1162
XP_005087312.1/1-1163
XP_003896906.1/1-1159
XP_014990765.2/1-1159
XP_014201355.2/1-1159
NP_000229.1/1-1159
NP_001180587.1/1-1158
XP_012641601.1/1-1160
XP_017375990.1/1-1159
JAB18309.1/1-1160
XP_023106495.1/1-1167
XP_040320396.1/1-1157
XP_005326715.1/1-1159
XP_036102470.1/1-1159
KAF6086072.1/1-1165
XP_045877243.1/1-1158
NP_001166444.1/1-1158
NP_001092571.1/1-849
1310
1320
1330
1340

**Fig. S1 – Multiple sequence alignment of ERG1 and EAG1 in mammal representative species.** 20 sequences of mammalian ERG1a and 20 sequences of mammalian EAG1 were used for the alignment. Gene access numbers are stated to the left, and the amino acid position within the alignment is in the central superior region. Residues in the alignment are colored according to the Kyte-Doolittle hydrophobicity scale available in Jalview (Kyte and Doolittle, 1982; Waterhouse et al., 2009). The most hydrophobic residues are colored red, and the most hydrophilic ones are colored blue. The colors in intermediate hydrophobicity residues vary from shades of purple according to their position in the scale.

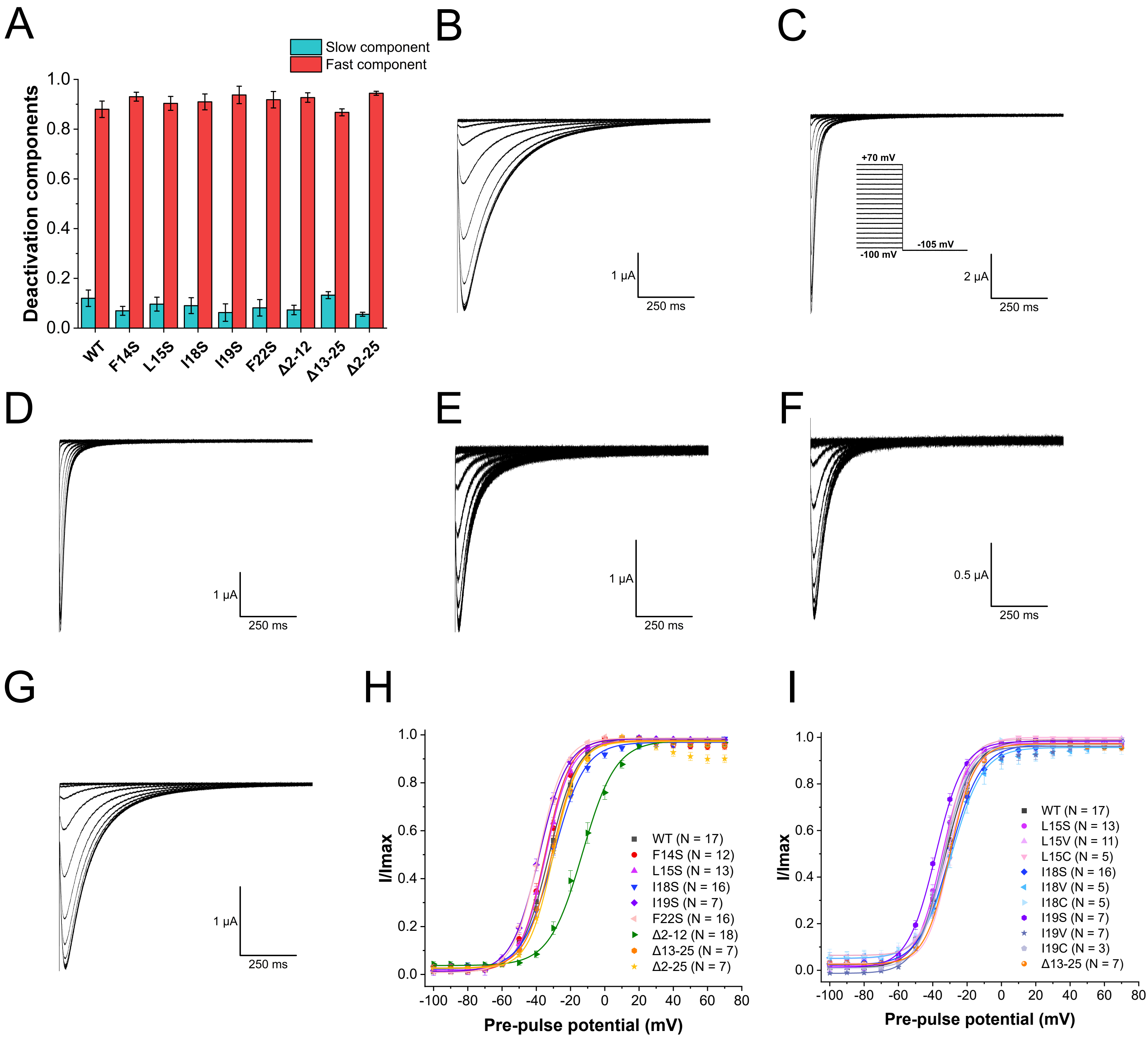
A
Slow component
Fast component
Deactivation components
1.0
0.8
0.6
0.4
0.2
0.0
WT
F14S
L15S
I18S
I19S
F22S
Δ2-12
Δ13-25
Δ2-25
B
1 μA
250 ms
C
+70 mV
-100 mV
-105 mV
2 μA
250 ms
D
1 μA
250 ms
E
1 μA
250 ms
F
0.5 μA
250 ms
G
1 μA
250 ms
H
I/Imax
WT (N = 17)
F14S (N = 12)
L15S (N = 13)
I18S (N = 16)
I19S (N = 7)
F22S (N = 16)
Δ2-12 (N = 18)
Δ13-25 (N = 7)
Δ2-25 (N = 7)
-100 -80 -60 -40 -20 0 20 40 60 80
Pre-pulse potential (mV)
I
I/Imax
WT (N = 17)
L15S (N = 13)
L15V (N = 11)
L15C (N = 5)
I18S (N = 16)
I18V (N = 5)
I18C (N = 5)
I19S (N = 7)
I19V (N = 7)
I19C (N = 3)
Δ13-25 (N = 7)
-100 -80 -60 -40 -20 0 20 40 60 80
Pre-pulse potential (mV)

**Fig. S2 – Analysis of the deactivation weights and the voltage dependences of activation. (A)** Analysis of the deactivation components. Relative amplitudes of the fast and slow components were calculated using the equation $A_f$ / ($A_f$ + $A_s$) and are reported in **Supplemental Table 2.1**. The fast component of deactivation has a major contribution at the applied voltage. Tail currents were evoked at -120 mV after a step to +60 mV (see *Materials and Methods* section). **(B-G)** Representative tail currents of WT **(B)** Δ2-12 **(C)**, Δ2-25, **(D)**, Δ13-25 **(E)**, I19S **(F)**, and F14S **(G)** hERG channels. **(H)** Analysis of the voltage dependence of activation of serine mutations and truncation of the PAS-cap α-helix. **(I)** Analysis of the voltage dependence of activation of cysteine and valine mutations of hydrophobic residues located at the PAS-cap α-helix and deletion of the PAS-cap α-helix. Tail currents were normalized as described in Materials and methods to produce I/Imax. Data were fitted with a Boltzmann function (curves). Data are mean ± SEM.

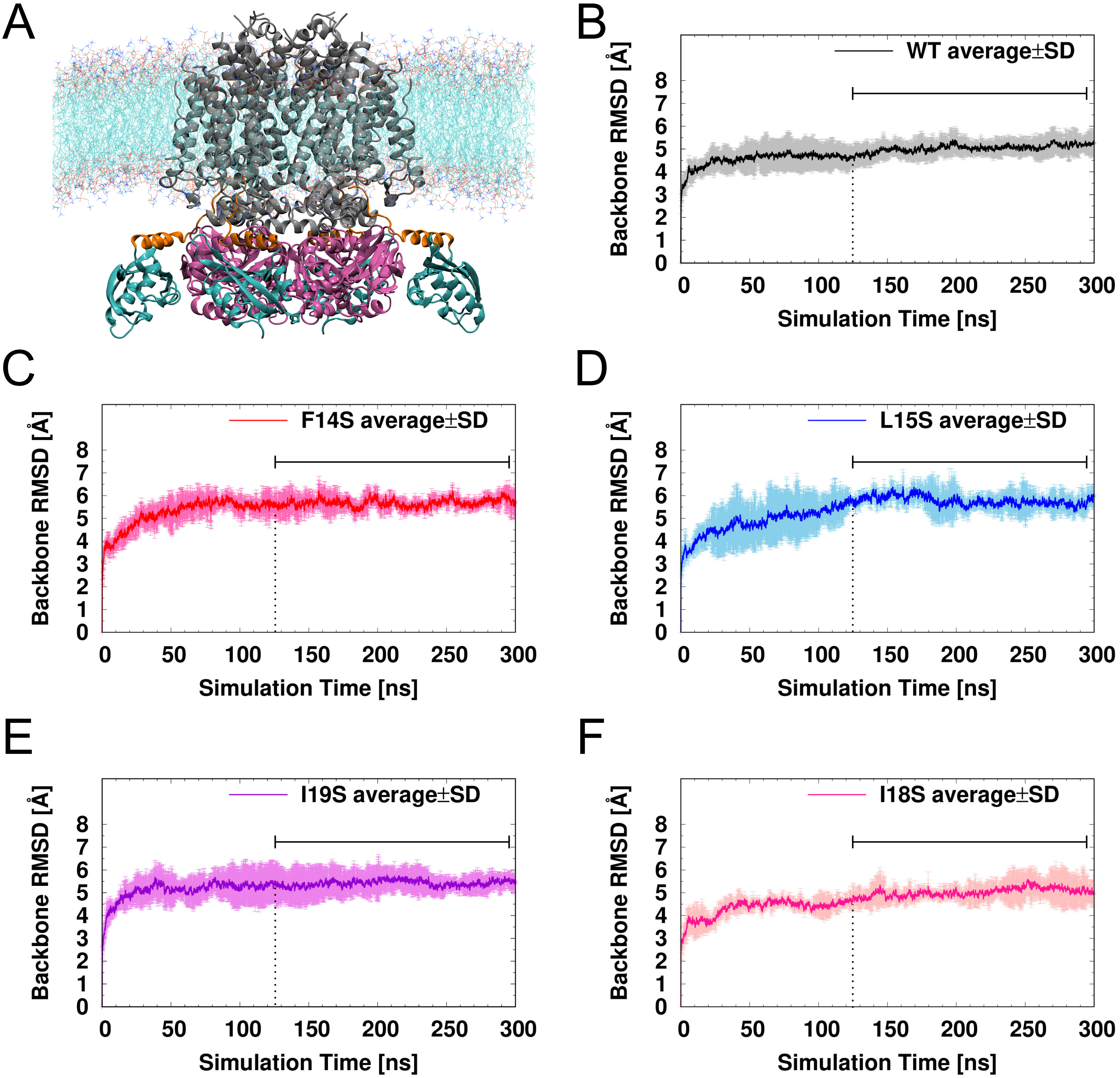
A
B
WT average±SD
Backbone RMSD [Å]
Simulation Time [ns]
C
F14S average±SD
Backbone RMSD [Å]
Simulation Time [ns]
D
L15S average±SD
Backbone RMSD [Å]
Simulation Time [ns]
E
I19S average±SD
Backbone RMSD [Å]
Simulation Time [ns]
F
I18S average±SD
Backbone RMSD [Å]
Simulation Time [ns]

**Fig. S3 – Equilibrium of the molecular dynamics simulations**. (**A**) Representation of the MDS system showing the hERG1a embedded in a POPC lipid bilayer; for better visualization, waters (SPC), $K^+$, and $Cl^-$ ions are not shown. (**B**) Root mean squared distance (RMSD) of WT and mutant hERG1a channels along the 300 ns MDS. The vertical dotted line indicates the time at which simulations were considered at equilibrium, while the horizontal black line indicates the time frame used for the analyses.

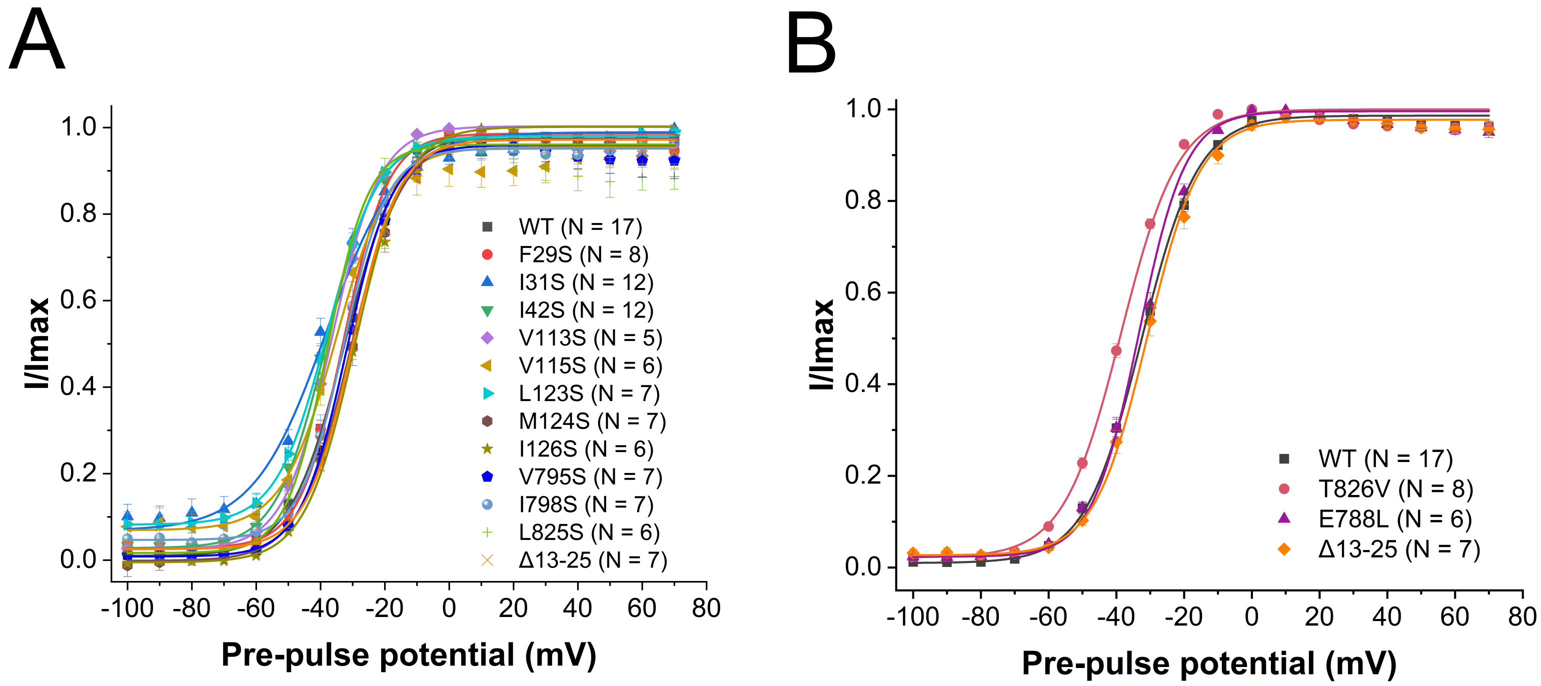
A
I/Imax
Pre-pulse potential (mV)
WT (N = 17)
F29S (N = 8)
I31S (N = 12)
I42S (N = 12)
V113S (N = 5)
V115S (N = 6)
L123S (N = 7)
M124S (N = 7)
I126S (N = 6)
V795S (N = 7)
I798S (N = 7)
L825S (N = 6)
Δ13-25 (N = 7)
B
I/Imax
Pre-pulse potential (mV)
WT (N = 17)
T826V (N = 8)
E788L (N = 6)
Δ13-25 (N = 7)

**Fig. S4 – Analysis of the voltage dependence of activation of mutations at the globular PAS and CNBhD.** I/Imax of WT and mutant hERG1a channels with serine mutations of hydrophobic residues at the globular PAS and CNBh domains. Tail currents were normalized as described in Materials and methods to produce I/Imax. Data were fitted with a Boltzmann function (curves). Data are mean ± SEM.

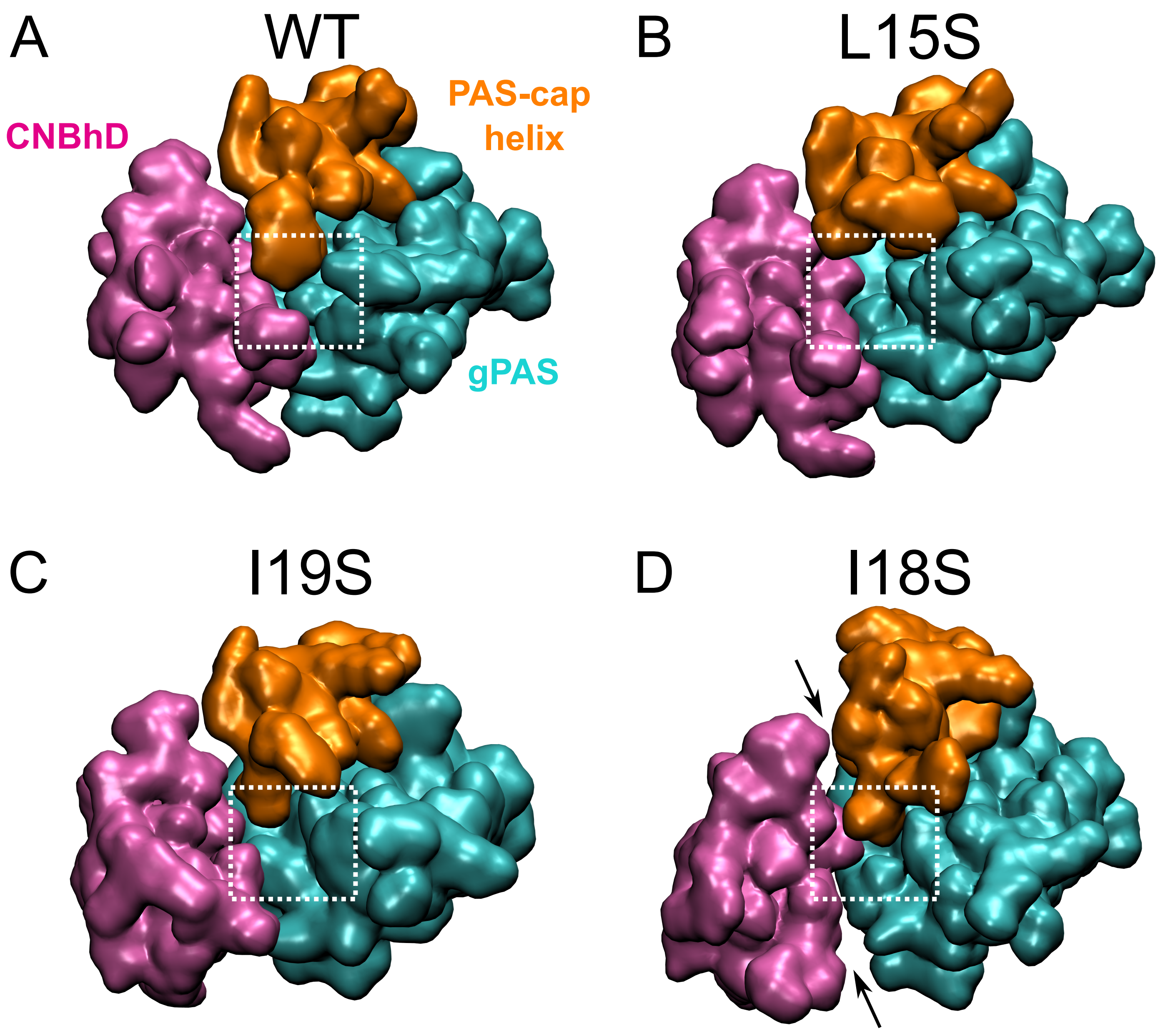
A
WT
PAS-cap
helix
CNBhD
gPAS
B
L15S
C
I19S
D
I18S

**Fig S. 5 – Serine substitutions of residues at the PAS-cap helix hydrophobic patch disrupt the tight interface of the hydrophobic nexus**. Conformation of the three-dimensional nexus in the WT **(A)**, L15S **(B)**, I19S **(C),** and I18S **(D)** channels. White squares represent the region forming the hydrophobic nexus. Black arrows in panel D highlight the loosened interface in the I18S mutant.

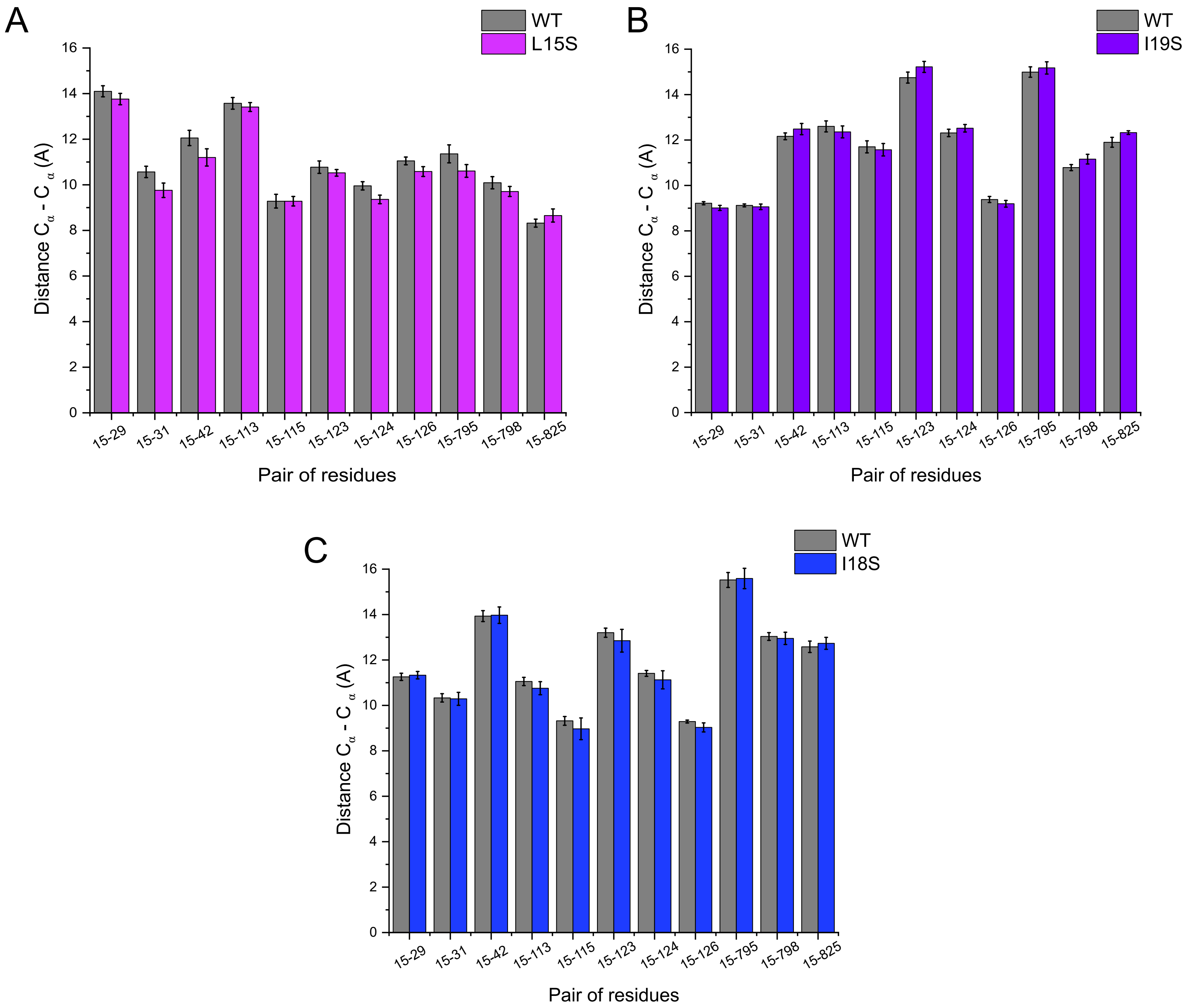
A
WT
L15S
Distance Cα - Cα (A)
Pair of residues
B
WT
I19S
C
WT
I18S
15-29
15-31
15-42
15-113
15-115
15-123
15-124
15-126
15-795
15-798
15-825
0
2
4
6
8
10
12
14
16

**Fig S. 6 – Serine substitutions of residues at the PAS-cap helix hydrophobic do not cause backbone movements of the implicated domains.** Comparison of the average distance between the Carbon α of residues contributing to the hydrophobic nexus in WT versus L15S (**A**), I19S (**B**), and I18S (**C**). Bars represent the average distance of the four subunits in three replicates ± S.E.M.

**Table S1 – Deactivation weights.** Relative amplitudes of the fast and slow components are shown ($A_f$ / ($A_f$ + $A_s$)). Note that deactivation is dominated by the fast component at -120 mV.

| **Mutant** | **$A_f$ / ($A_f$ + $A_s$), $_{-120mV}$** | **N** |
|---|---|---|
| WT | 0.88 ± 0.00 | 18 |
| F14S | 0.93 ± 0.00 | 12 |
| L15S | 0.90 ± 0.00 | 6 |
| I18S | 0.91 ± 0.00 | 18 |
| I19S | 0.93 ± 0.01 | 5 |
| F22S | 0.90 ± 0.00 | 9 |
| Δ2-12 | 0.93 ± 0.00 | 15 |
| Δ13-25 | 0.87 ± 0.00 | 10 |
| Δ2-25 | 0.94 ± 0.00 | 11 |

**Table S2 – Current decay time constants.** Deactivation time constants of currents recorded at −120 mV from a +60 mV pre-pulse in *Xenopus laevis* oocytes expressing WT, deletion, and PAS cap α helix mutant channels. The decay of current at -120 mV was fitted with a double exponential function to obtain fast ($\tau_{fast,\ -120\ mV}$) and slow ($\tau_{slow,\ -120\ mV}$) time constants for deactivation.

| **Mutant** | $\tau_{fast,\ -120mV}$ **(ms)** | $\tau_{slow,\ -120mV}$ **(ms)** | **N** |
|---|---|---|---|
| WT | 80.04 ± 1.55 | 486.05 ± 28.65 | 18 |
| F14S | 71.81 ± 1.76 | 523.49 ± 22.47 | 12 |
| L15S | 27.74 ± 0.56 | 208.24 ± 1.51 | 6 |
| I18S | 32.05 ± 1.37 | 266.62 ± 5.45 | 18 |
| I19S | 40.86 ± 0.51 | 347.00 ± 8.45 | 5 |
| F22S | 88.66 ± 2.36 | 618.58 ± 14.76 | 9 |
| Δ2-12 | 18.58 ± 0.90 | 261.42 ± 3.30 | 15 |
| Δ13-25 | 38.26 ± 0.78 | 230.55 ± 4.37 | 10 |
| Δ2-25 | 17.23 ± 0.36 | 166.40 ± 2.19 | 11 |
| L15V | 86.05 ± 1.11 | 370.61 ± 4.25 | 12 |
| L15C | 84.66 ± 1.37 | 427.18 ± 6.59 | 9 |
| I18V | 58.80 ± 0.40 | 339.27 ± 4.67 | 9 |
| I18C | 63.30 ± 1.53 | 489.39 ± 10.64 | 9 |
| I19V | 79.64 ± 1.35 | 539.12 ± 8.72 | 9 |
| I19C | 83.23 ± 2.22 | 590.59 ± 20.06 | 8 |

**Table S3 – Voltage dependence of activation of WT, deletion, and PAS-cap α-helix mutant hERG1a channels.** Boltzmann parameters were obtained by fitting IV plot for hERG tail currents to a Boltzmann equation (see Methods) using Origin 2021b (OriginLab). All data are means ± SE. $V_{1/2}$, voltage at which half of the maximal current is achieved. k, is the slope factor of the I-V relation. N is the number of replicates.

| Condition | $V_{1/2}$ (mV) | k (mV) | N |
|---|---|---|---|
| WT | -32.56 ± 0.57 | 8.37 ± 0.34 | 17 |
| F14S | -34.96± 1.18 | 7.41 ± 0.64 | 12 |
| L15S | -34.52 ± 0.29 | 7.45 ± 0.20 | 13 |
| I18S | -30.03 ± 0.63 | 9.04 ± 0.46 | 16 |
| I19S | -38.30 ± 0.35 | 8.12 ± 0.24 | 7 |
| F22S | -38.70 ± 0.72 | 7.47 ± 0.46 | 16 |
| Δ2-12 | -13.04 ± 0.68 | 9.69 ± 0.39 | 18 |
| Δ13-25 | -30.92 ± 0.69 | 7.78 ± 0.42 | 7 |
| Δ2-25 | -30.04 ± 1.25 | 7.28 ± 0.72 | 7 |
| L15V | -30.80 ± 0.86 | 8.39 ± 0.59 | 11 |
| L15C | -29.47 ± 1.26 | 6.74 ± 0.56 | 5 |
| I18V | -28.79 ± 0.92 | 9.00 ± 0.69 | 5 |
| I18C | -32.72 ± 1.32 | 6.67 ± 0.80 | 5 |
| I19V | -34.09 ± 0.70 | 7.77 ± 0.41 | 7 |
| I19C | -33.31 ± 0.65 | 7.19 ± 0.24 | 3 |

**Table S4 – Current decay time constants.** Deactivation time constants of currents recorded at −120 mV from a +60 mV pre-pulse in *Xenopus laevis* oocytes expressing WT, deletion, globular PAS and CNBhD mutant channels. The decay of current at -120 mV was fitted with a double exponential function to obtain fast ($\tau_{fast,\ -120\ mV}$) and slow ($\tau_{slow,\ -120\ mV}$) time constants for deactivation.

| **Mutant** | $\tau_{fast,\ -120mV}$ **(ms)** | $\tau_{slow,\ -120mV}$ **(ms)** | **N** |
|---|---|---|---|
| WT | 80.04 ± 1.55 | 486.05 ± 28.65 | 18 |
| F29S | 48.60 ± 1.54 | 323.53 ± 5.30 | 9 |
| I31S | 36.56 ± 2.26 | 283.91 ± 5.18 | 12 |
| I42S | 52.85 ± 1.24 | 383.86 ± 5.20 | 12 |
| V113S | 62.47 ± 0.97 | 327.87 ± 3.20 | 7 |
| V115S | 44.67 ± 0.70 | 291.04 ± 6.05 | 13 |
| L123S | 66.73 ± 1.09 | 336.33 ± 5.26 | 15 |
| M124S | 82.49 ± 1.52 | 375.95 ± 4.19 | 22 |
| I126S | 59.74 ± 3.49 | 343.90 ± 7.64 | 6 |
| V795S | 44.07 ± 0.63 | 338.29 ± 6.11 | 9 |
| I798S | 25.40 ± 0.87 | 233.26 ± 2.50 | 7 |
| L825S | 31.63 ± 0.73 | 250.18 ± 2.21 | 8 |
| Δ13-25 | 38.26 ± 0.78 | 230.55 ± 4.37 | 10 |

**Table S5 – Voltage dependence of activation of hERG1a globular PAS and CNBh domain mutations.** Boltzmann parameters were obtained by fitting IV plot for hERG tail currents to a Boltzmann equation (see Methods) using Origin 2021b (OriginLab). All data are means ± SE. $V_{1/2}$, voltage at which half of the maximal current is achieved. k, is the slope factor of the I-V relation. N is the number of replicates.

| Condition | $V_{1/2}$ (mV) | k (mV) | N |
|---|---|---|---|
| WT | -32.56 ± 0.57 | 8.37 ± 0.34 | 17 |
| F29S | -32.87 ± 1.07 | 6.87 ± 0.70 | 8 |
| I31S | -38.17 ± 2.31 | 11.28 ± 1.47 | 12 |
| I42S | -38.20 ± 0.35 | 7.82 ± 0.21 | 12 |
| V113S | -36.46 ± 0.56 | 7.47 ± 0.43 | 5 |
| V115S | -35.18 ± 1.29 | 8.36 ± 1.08 | 6 |
| L123S | -37.72 ± 0.33 | 8.00 ± 0.24 | 7 |
| M124S | -30.72 ± 0.57 | 7.99 ± 0.40 | 7 |
| I126S | -29.32 ± 1.16 | 7.80 ± 0.79 | 6 |
| V795S | -31.96 ± 0.48 | 7.15 ± 0.28 | 7 |
| I798S | -32.77 ± 0.41 | 6.82 ± 0.28 | 7 |
| L825S | -37.89 ± 0.10 | 6.44 ± 0.61 | 6 |
| Δ13-25 | -30.92 ± 0.69 | 7.78 ± 0.42 | 7 |
